# Structural variant selection for high-altitude adaptation using single-molecule long-read sequencing

**DOI:** 10.1101/2021.03.27.436702

**Authors:** Jinlong Shi, Zhilong Jia, Xiaojing Zhao, Jinxiu Sun, Fan Liang, Minsung Park, Chenghui Zhao, Xiaoreng Wang, Qi Chen, Xinyu Song, Kang Yu, Qian Jia, Depeng Wang, Yuhui Xiao, Yinzhe Liu, Shijing Wu, Qin Zhong, Jue Wu, Saijia Cui, Xiaochen Bo, Zhenzhou Wu, Manolis Kellis, Kunlun He

**Affiliations:** Key Laboratory of Biomedical Engineering and Translational Medicine, Ministry of Industry and Information Technology, Chinese PLA General Hospital, Beijing, China; Beijing Key Laboratory for Precision Medicine of Chronic Heart Failure, Chinese PLA General Hospital, Beijing, China; Research Center of Medical Artificial Intelligence, Chinese PLA General Hospital, Beijing, China; Research Center of Medical Big Data, Chinese PLA General Hospital, Beijing, China; GrandOmics Biosciences Inc, Beijing, China; Beijing Institute of Radiation Medicine, Beijing, China; BioMind Inc, Beijing, China; Computer Science and Artificial Intelligence Laboratory, Massachusetts Institute of Technology, Cambridge, MA, USA; Broad Institute of MIT and Harvard, Cambridge, MA, USA

## Abstract

Structural variants (SVs) can be important drivers of human adaptation with strong effects, but previous studies have focused primarily on common variants with weak effects. Here, we used large-scale single-molecule long-read sequencing of 320 Tibetan and Han samples, to show that SVs are key drivers of selection under high-altitude adaptation. We expand the landscape of global SVs, apply robust models of selection and population differentiation combining SVs, SNPs and InDels, and use epigenomic analyses to predict driver enhancers, target genes, upstream regulators, and biological functions, which we validate using enhancer reporter and DNA pull-down assays. We reveal diverse Tibetan-specific SVs affecting the cis- and trans-regulatory circuitry of diverse biological functions, including hypoxia response, energy metabolism, lung function, etc. Our study greatly expands the global SV landscape, reveals the central role of gene-regulatory circuitry rewiring in human adaptation, and illustrates the diverse functional roles that SVs can play in human biology.

## Introduction

Structural variants (SVs) account for the majority of variable base pairs in the human genome and can cause dramatic alterations in gene function and gene regulation. They have been shown to play important biological roles in human biology and human disease^1^. For example, an inversion disconnecting TFAP2A from its enhancers causes branchiooculofacial syndrome^2^, and copy number differences in the AMY1 gene are associated with high-starch diets or low-starch diets in the population^3^. Despite such dramatic examples, the roles of SVs in disease and in human evolutionary adaptation remain poorly studied, with most studies focusing instead on common single-nucleotide variants of weak effect, due in great part to technological limitations.

The adaptation of Tibetan people to high altitude provides an ideal model for studying adaptation during the evolutionary history of modern humans given its well-controlled context^4–7^, but this adaptation remains insufficiently studied at the population scale. Altitude sickness mainly takes the form of acute mountain sickness, high-altitude pulmonary oedema, and high-altitude cerebral oedema, which involve dizziness, headache, muscle aches, etc. The prevalence rate of these types of altitude sickness in the Tibetan population is lower than that in the Han population. Previous studies^4,8^ have focused mainly on hypoxia-inducible factor (HIF) pathways, including those involving *EGLN1* and *EPAS1*, the latter of which shows a Denisovan-like haplotype in Tibetans^6^. However, these early studies relied on small sample sizes that were unlikely to reveal population genetic characteristics, used short-read next-generation sequencing (NGS) techniques that are not well suited for SV analysis, and lacked the sizable control cohorts of Han populations^9^ necessary for revealing the unique genetic characteristics of Tibetan adaptation.

By contrast, long-read technologies, including single-molecule real-time (SMRT) and Oxford Nanopore Technologies (ONT) platforms, can provide a complete view of genomic variation. SVs can be important drivers of genome biological function and human evolutionary adaptation by enabling the rewiring of the long-range gene regulatory circuitry, amplification of gene clusters, and strong-effect adaptive changes that can involve multiple genes. Even in very small sample sizes, long-read sequencing has revealed extensive variation in SVs in the human population^10^ and revealed that transposable elements, including long interspersed nuclear elements (LINEs) and short interspersed nuclear elements (SINEs) can be used to recapitulate the patterns of human evolution^11^, underlie most reported SVs^10^, and contribute to both medically important and evolutionarily selected variation^12^. However, the pattern of SV hotspots in the human genome remains incompletely understood, and comprehensive studies of large cohorts are needed to understand the role of SVs in human adaptation.

Here, we used long-read sequencing technologies to evaluate the roles of SVs in recent human adaptation (**Fig. 1a**). Our study reveals the first (unique) SV landscape for ethnic Han and Tibetan populations at a large scale. This work provides a large call set of 122,876 total SVs and contributes 102,103 novel SVs to the existing SV call set for East Asians. Unique patterns of SVs are also explored, which provides a different perspective for understanding genome evolution. Further comparisons of population stratification elucidate the comprehensive genetic landscape of the Han and Tibetan populations, revealing the potential roles of SVs in evolutionary adaptation to the high altitudes inhabited by Tibetans. We provide high-altitude adaptation candidate SVs, showing their functional impacts on enhancers, exons and the 3D genome. We show the functional connections between SV-associated genes and the unique traits of Tibetans. These findings implicate multiple genes in biologically relevant pathways, including the HIF, insulin receptor signalling, inflammation, and glucose and lipid metabolism pathways. Moreover, experimental validation confirms our analytical result showing that the most Tibetan-specific SV, a deletion downstream of EPAS1, disrupts the super enhancer in this genomic area in Tibetans and affects the binding of regulatory molecules critical for gene transcription activities. Our study expands the known East Asian and global SV sets and highlights the functional impacts and adaptation of SVs in Tibetans via complex cis- and trans-regulatory circuitry rewiring.

**Figure 1.**
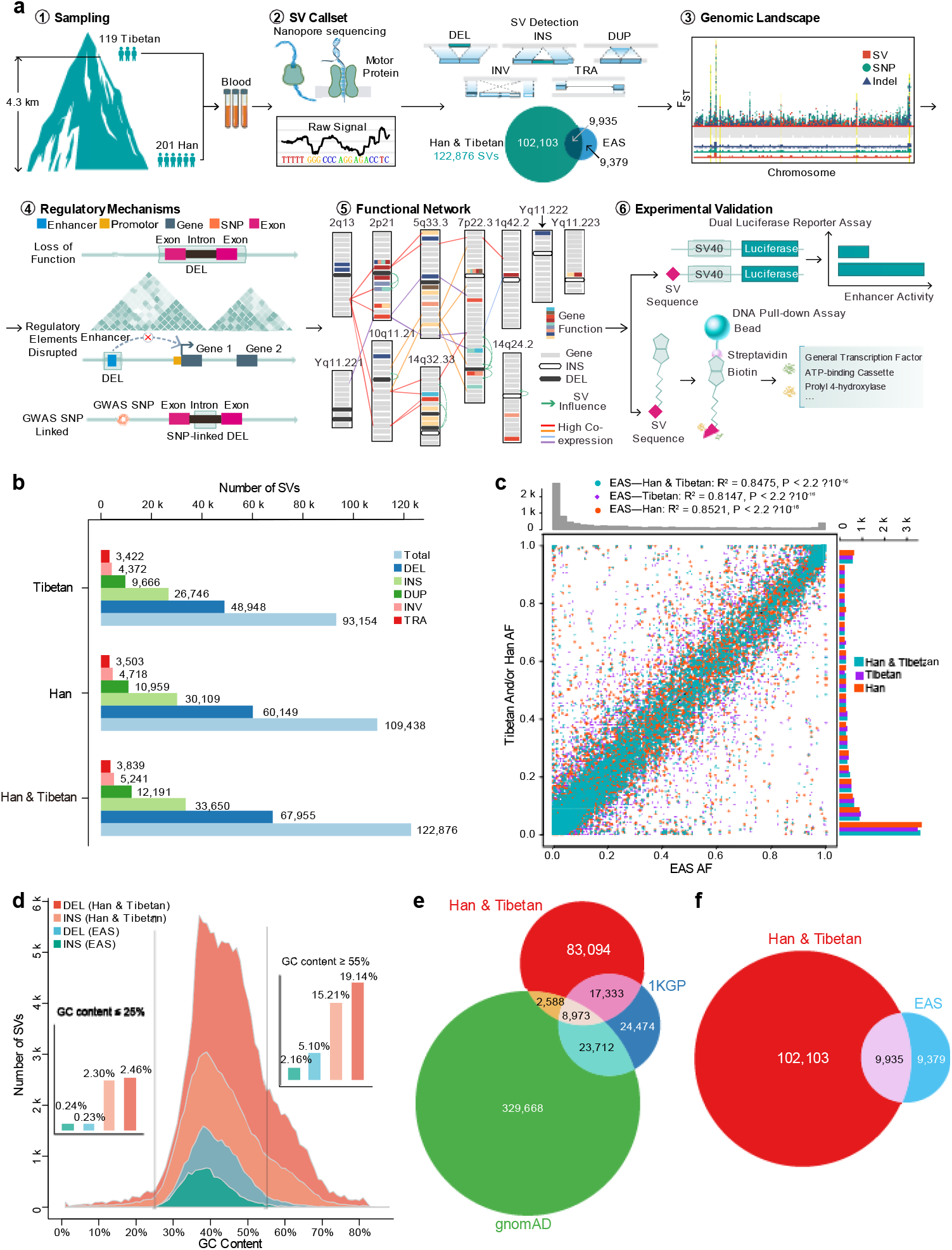
SV discovery in 119 Tibetan and 201 Han samples. **a.** Summary of the experimental pipeline. Overall, 320 Han and Tibetan samples were collected and sequenced via the ONT platform, resulting in 122,876 SVs. Candidate SVs for high-altitude adaptation, their functional regulatory mechanisms based on their connections with exons, enhancers and TAD boundaries, and candidate genes for high-altitude adaptation were explored. Cis- and trans-regulatory circuitry rewiring was validated in two biological assays. **b.** The numbers of 5 types of SVs in Han, Tibetan, and all samples. The majority are deletions and insertions. **c.** Allele frequency consistency between the SVs in the Han and Tibetan cohort and the 1KGP EAS cohort. The high consistency between them indicates the high quality of our SV call set. **d.** GC content distribution of insertions and deletions in the Han and Tibetan cohort and the EAS cohort, indicating that the ONT platform performs well even in genomic regions with a biased GC content. **e.** Overlap between the SVs of our cohort and 1KGP and gnomAD cohorts, showing 83,094 novel SVs. **f.** Overlap between the SVs of our cohort and the 1KGP EAS cohort, showing a 6-fold increase in the number of previously annotated East Asian SVs.

## Results

### 1. Constructing and validating the de novo SV call set

#### SV discovery and quality evaluation

We sequenced the genomes of 201 Han and 119 Tibetan individuals using the ONT PromethION platform with an average depth of 21-fold coverage (**Supplementary Table 1a**). We detected 109,418 SVs in the Han population and 93,154 SVs in the Tibetan population (**Fig. 1b**), resulting in 122,876 SVs in total (89% and 76%, respectively). These SVs consisted of 67,955 deletions, 33,650 insertions, 12,191 duplications, 5,241 inversions, and 3,839 translocations. Each sample, averagely contains 20,741 SVs, including 8,520 insertions, 9,392 deletions, 1,908 duplications, 548 inversions and 373 translocations (**Supplementary Fig. 1a** and **Supplementary Table 1a**). The numbers of different types of SVs showed no large difference in each Han and Tibetan sample (**Supplementary Fig. 1a**). Remarkably, in 50% of our samples, more than 90% of the SVs in the Han and Tibetan populations were captured, indicating that our sample size was sufficient to comprehensively profile the SV landscape of the Han and Tibetan populations (**Supplementary Fig. 1b-d**).

We used several lines of evidence to confirm the high quality of our Han and Tibetan SV call set. First, the manual curation of 298 SVs across all samples showed 94% accuracy (**Supplementary Table 1b**). Second, the PCR validation of 4 SVs in 57 samples showed 96% accuracy (**Supplementary Table 1c**). Third, our SV allele frequencies showed a Pearson correlation of 0.92 with the East Asian (EAS) database of the 1000 Genomes Project (1KGP) phase 3 (**Fig. 1c**). Fourth, 74% (13,342) of the SVs were shared between the sequencing results obtained for one sample using both the ONT (12.2X depth, 36.59 Gb, 18,002 SVs) and SMRT circular consensus sequencing (15.4X depth, 452 Gb, 20,617 SVs) platforms (**Supplementary Tables 1d, 2, 3**)^13^. Notably, 71% (12,429) of the SVs were shown to be shared when the new version of Guppy (3.0.5) was used (**Supplementary Table 3**), indicating no significant change in the overall quality of the SV call sets. These results collectively indicate the high quality of the SV call set. Accordingly, we provide an authoritative new reference set for future studies of genetic variation.

We also confirmed that the long-read sequencing platform performed well even in genomic regions with a GC-biased base composition. We compared the GC composition of deletions and insertions, two major types of SV, identified in our Han and Tibetan cohort and the EAS population of the 1000 Genomes database. The ONT platform revealed more SVs in total and more SVs in GC-biased areas than with NGS in the EAS population. For example, among all 67,954 deletions discovered using the ONT platform, 19.14% of the deletions were located in the high-GC content (≥55%) areas. Among all 12,969 deletions discovered using the NGS platform, only 5.10% of deletions were identified in high-GC content areas (**Fig. 1d**). This indicates the advantages of long-read sequencing for calling SVs, especially in genomic regions with a GC-biased base composition.

#### Comparison with existing SV call sets

We contribute a large number of new SVs and provide a useful reference panel for Chinese, East Asian, and worldwide populations. As an indicator of the near completeness of the SV results, we found that the majority of the identified SVs were shared between the Han and Tibetan populations and that the number of SVs in the Han population increased by only 17%, despite profiling almost twice as many Han as Tibetan genomes. We compared our SV set with two NGS-based SV call sets, those of the 1KGP^14^ and the Genome Aggregation Database (gnomAD)^15^. Approximately 28,894 SVs included in the 1KGP and gnomAD sets were reidentified in our call set (**Fig. 1e**). Importantly, our study expands the total number of SVs in global SV databases by ~37%, by adding 83,094 novel SVs, and increases the number of EAS SVs to 121,417, by contributing 102,103 novel SVs (**Fig. 1f**). The application of ONT sequencing clearly contributed greatly to the increase in the number of SVs.

Long-read sequencing of a large-scale sample of individuals is necessary for comprehensive population-scale SV profiling. We compared our SV set with an SV set derived from 15 samples using the SMRT platform^10^. Notably, 61,502 SVs (~61.7%) in our SV set were merged into 36,528 SVs (insertion length was ignored during SV merging due to in the absence of this information for the 15 genomes). Nevertheless, 74,138 novel SVs (**Supplementary Fig. 1e**) were identified compared with the SV set based on the 15 genomes. Furthermore, 59,878 novel SVs were found in our study (**Supplementary Fig. 1f**), the majority of which were low-frequency (allele frequency<0.1, ~58%) and singleton SVs (~29.9%) (**Supplementary Table 4**). We also compared our Tibetan SV set with the ZF1 SV call set, which was recently collected from a high-quality de novo assembled Tibetan genome based on the SMRT platform^16^. We found that only 15,890 SVs (~17%) in our Tibetan SV reference panel overlapped with the ZF1 SV call set (17,714 SVs) (**Supplementary Table 5**). A number of SVs overlapping with other publicly available SV sets verified the high quality of our Han and Tibetan SV set from another perspective. Thus, we provide a high-quality SV call set for a large-scale Han and Tibetan cohort.

### 2. Genome-wide properties of SVs

#### The SV landscape in the Han-Tibetan population

More SVs are distributed in repeat regions, such as regions containing transposable elements and satellite repeat regions. SVs showed 4-fold enrichment in repeat elements (81% occurred in repeats), including 40% of SVs in transposable elements (23% SINEs, 11% LINEs, 6% LTRs) and 27% in satellite repeat regions (22% in simple repeats, 5% satellite repeats) (**Fig. 2a**). SVs were 1.8-fold enriched in SINEs (23% vs. 13% expected) but 1.9-fold depleted in LINEs (11% vs. 22% expected) (**Fig. 2a**), possibly due to the increased selective pressure related to their longer length^17^. Within different allele frequency intervals, more SVs were distributed in repeat regions (**Supplementary Table 6**). These repeats are more likely to produce SVs because repeats are prone to non-allelic homologous recombination, replication slippage, and non-homologous end-joining^18^.

**Figure 2.**
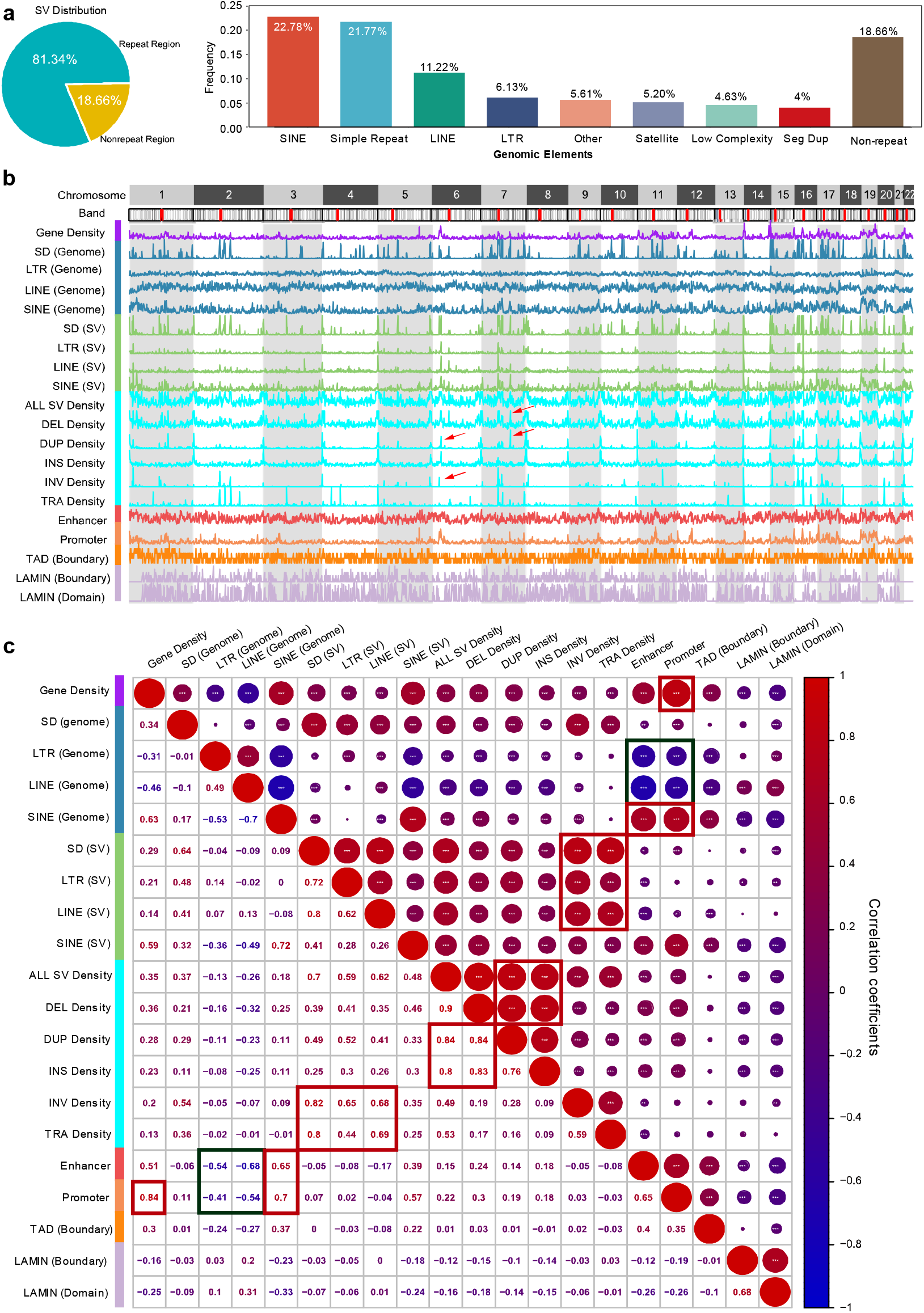
SV composition, length frequency and chromosome distributions. **a.** SV proportions in different genome regions. The majority of SVs are associated within repeat elements, such as LINEs and SINEs as shown in the pie chart. **b.** Densities of repeats and their-associated SVs, all SVs, regulatory elements and genes in the genome. There are hotspots (peaks with red arrow) of SVs in the genome. **c.** Correlations between repeat elements, their-associated SVs, regulatory elements and genes in the genome. Different types of SVs are highly positively (red) inter-correlated. High correlation coefficients are boxed (red for positive and blue for negative).

The different types of SVs are distributed among different types of repeat and functional elements in a biased manner. The enrichment among these elements differed among different SV types, with deletions, insertions, and duplications being associated with SINEs, LINEs, and simple repeats and inversions and translocations being associated with LINEs and satellite repeats (**Supplementary Table 7**). The SV length distribution showed two distinct peaks, at ~300 bp (Alu) and ~6 kb (LINE) (**Supplementary Fig. 2a, 2b**). Exonic SVs showed 6-fold enrichment relative to the exons in the genome (**Supplementary Fig. 2c**, **Supplementary Table 8**). LINE-associated SVs showed 2-fold depletion in intronic regions and a 5-fold enrichment in exonic regions (**Supplementary Fig. 2c**, **SupplementaryTable 8**). Concerning the SINE- and LINE-mediated SVs, duplications and inversions were >5-fold enriched in exons, while deletions were 40% depleted and insertions were 30-fold depleted in exons (**Supplementary Fig. 2d**, **Supplementary Table 8**), suggesting deleterious effects of SINE- and LINE-mediated insertions on the genome overall. The majority of SVs (75% SVs) occur at a low frequency (7.8%), while a greater number of higher-frequency insertions are maintained relative to other types of SVs (**Supplementary Fig. 2e**).

#### SV hotspots in the genome

SV hotspots in the genome were indicated by the extremely high density of SVs in certain regions relative to other regions in the whole genome (**Fig. 2b**). For example, there were 58 C7orf50-associated SVs, indicating that this region is prone to DNA breaks and the formation of SVs (**Supplementary Fig. 2f**). Deletions, insertions, and duplications showed a highly inter-correlated distribution in the genome (**Supplementary Fig. 2c**). The distributions of LINE-associated SVs, SD-associated SVs and translocations were also highly inter-correlated, whereas the distributions of LINEs and SDs in the human genome were anti-correlated. These results revealed clear SV hotspots in the genome. As promoters and enhancers are correlated with SINEs, they are also correlated with SINE-associated SVs. Notably, the distributions of promoters and enhancers were correlated with those of deletions, duplications, insertions, SINE-related SVs and SINEs but anti-correlated with those of LINEs and LTRs in the genome. This indicates that SINE-associated SVs, deletions, duplications, and insertions play more important roles in regulating gene transcription than other types of SVs.

#### Preferential evolutionary selection for SVs

We also found that SVs were very infrequent in exonic regions (17%), upstream regions (0.5%), 3’-UTRs (0.34%), and 5’-UTRs (0.16%), where they are more likely to disrupt functional elements, but that they were very abundant in intronic regions (30%) and intergenic regions (46%), where they are less likely to be disruptive (**Supplementary Fig. 2c**). This difference was also observed in the fraction of fully penetrant (allele frequency=1 within the corresponding population) SVs vs. singleton SVs in each population, with a larger fraction of exonic SVs being singletons and a larger fraction of intronic and intergenic SVs being shared, consistent with continued evolutionary pressures acting on SV allele frequency in each population (**Supplementary Fig. 2h** and **Supplementary Table 1e**).

Genome evolution selects and preserves more repeat-associated SVs. We found that in both the Han and Tibetan populations, fully penetrant SVs were ~6.6 times more frequent in repeat regions than in non-repeat regions, while singleton SVs (found in only one individual in a given population) were only ~3.8 times more frequent in repeat regions than in non-repeat regions (**Supplementary Fig. 2g**). This indicates that SVs are preferentially excluded (singletons) from non-repeat regions, where they are more likely to disrupt functional elements, and that they are preferentially tolerated (high allele frequency) in repeat elements, where they accumulate over evolutionary time.

We also found that while deletions and insertions were similarly abundant in the genome, deletions were more likely to be singletons than to be shared, while insertions were more likely to be shared than to be singletons (**Supplementary Fig. 2i, SupplementaryTable 1e**), indicating that deletions are more likely to disrupt elements than insertions.

In summary, our comparisons of the frequency of SVs in different genomic regions and of the relative frequencies of singletons and shared SVs indicate the preferential retention of repeat-associated SVs, intronic SVs and insertions vs. deletions, suggesting that these variant types are less likely to disrupt genomic functions.

### 3. Population genetics of Han-Tibetan populations and the role of functional SVs in evolutionary adaptation

#### SVs are a representative ethnic characteristic

The Han and Tibetan SV call sets provide an unprecedented resource for the in-depth analysis of genomic variations for comparisons between Han and Tibetan populations. The Han and Tibetan cohorts shared 79,716 SVs (**Supplementary Fig. 3a**). A principal component analysis (PCA) established that SVs could be used to clearly distinguish these two very closely related populations (**Supplementary Fig. 3b**). These results suggested the existence of significant genetic differences between the Han and Tibetan populations based on SVs alone, which are usually revealed by SNP analysis. When we extended the PCA to other populations from the 1KGP database, such as African, American, EAS, European, and South Asian populations, the Han and Tibetan populations were shown to be closely related to EAS populations and distant from these other populations (**Fig. 3a**). This indicates that the Tibetan population is genetically closer to the Han population than to other populations and that these populations probably originated from a single common ancestor. Admixture analysis using the SV call set clearly showed separation between the Han and Tibetan populations (**Fig. 3b**, top). The hierarchical clustering of the SVs with F_ST_ >0.25 also showed that the Han and Tibetan populations possess ethnicity-specific SVs as well as common SVs (**Fig. 3b**, bottom). Evolutionary tree analysis based on all the SVs also showed a clear separation between Han and Tibetan with 3 sample exception (**Fig. 3c**). These results demonstrate that SVs, in addition to SNPs, are a powerful proxy for distinguishing genetically closely related populations as a representative characteristic of an ethnic group.

**Figure 3.**
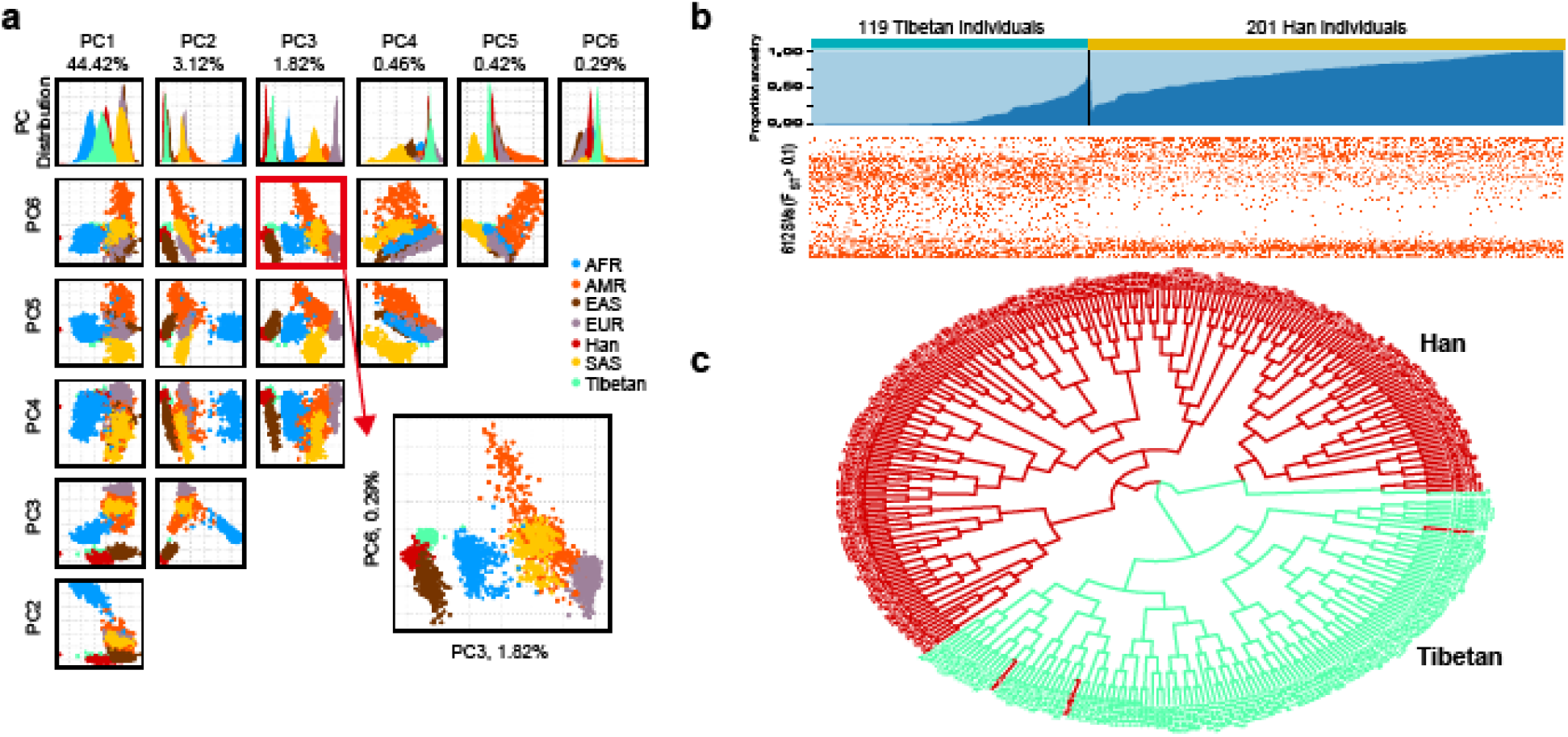
Population genetics of Han-Tibetan populations. **a.** PCA of the SV call sets of the Tibetan and Han cohorts and 1KGP African (AFR), American (AMR), East Asian (EAS), European (EUR) and South Asian (SAS) cohorts. The Han (red) and Tibetan (green) populations are close to the EAS (brown) populations, as expected, and can be clearly separated according to PC3 and PC6. **b.** Population structure of the Tibetan and Han populations. Admixture analysis (top), clustered SVs with an F_ST_>0.1 (bottom). The SVs can distinguish two populations, although the number of SVs per individual in the two populations is similar. **c.** A clear separation between Han and Tibetan with 3 sample exception in the evolutionary tree analysis based on all the SVs of Han (red) and Tibetan (green).

#### Genetic landscapes of Han-Tibetan populations

SVs, InDels, and SNPs function synergistically to allow Tibetans to live in high-altitude environments. We complemented our long-read sequencing results with deep short-read sequencing in 148 samples and carried out a comprehensive comparison of the Han-Tibetan genome based on population NGS/TGS sequencing data, comprehensively revealing the genomic signals of evolutionary selection for high-altitude adaptations. A Manhattan plot based on SNPs, InDels, and SVs between the Han and Tibetan populations revealed clear evolutionary selection (**Fig. 4a**). The selection for these 3 types of genetic variations was highly consistent in multiple genomic regions, and several regions were found to differ significantly between the Han and Tibetan populations (**Supplementary Fig. 4a**). For example, there are many SNPs, InDels, and SVs in the region around EGLN1 on chromosome 1 (**Fig. 4b**) and the region around EPAS1 on chromosome 2 (**Fig. 4c**), forming several plateaus of evolutionary selection in the Manhattan plot of the human genome.

**Figure 4.**
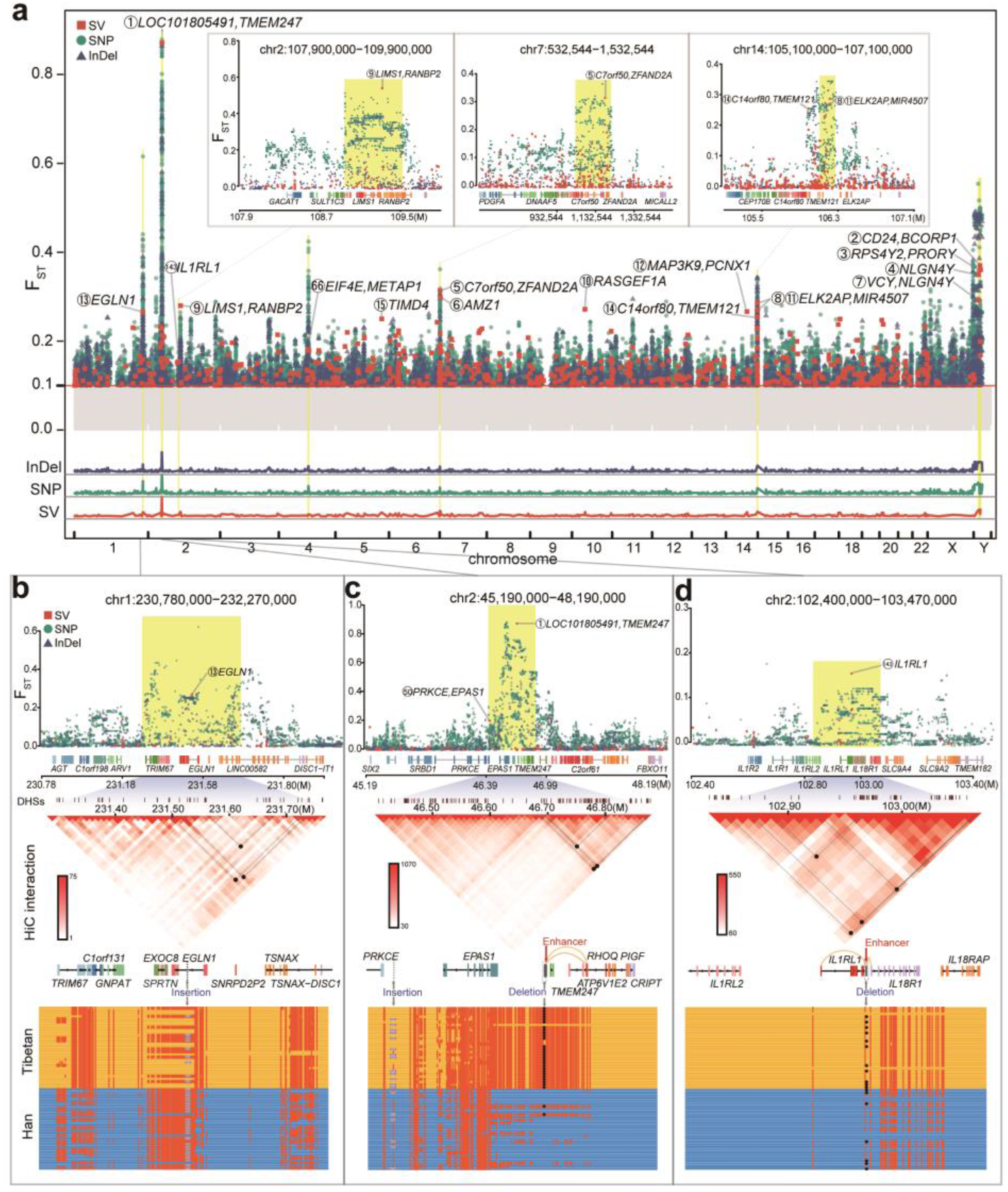
Comparison of Han-Tibetan populations reveals the genetic landscape of evolutionary adaptation. **a.** Manhattan plot (top) based on the FST values of SVs (orange-red boxes), SNPs (blue-green dots) and InDels (dark blue triangles) between the Han and Tibetan cohorts. Overall, 15 SVs with an F_ST_>0.25 are highlighted (yellow), and associated genes are labelled (black) with rank (circled number). The shadows in the Manhattan plot for SVs, SNPs and InDels (bottom) suggest the high consistency of evolutionary selection for SVs, SNPs and InDels. **b.** Manhattan plot (top), TAD (middle) and haplotype (bottom) near EGLN1 on chromosome 1 based on the FST values of SNPs, InDels, and SVs between the Han and Tibetan cohorts. A 131 bp insertion (grey) in the intron of EGLN1 shows high LD with SNPs (red dot) and potentially disrupts a loop (black dot and inverted triangle) boundary. **c.** Manhattan plot (top), TAD (middle) and haplotype (bottom) near EPAS1 and TMEM247 on chromosome 2 based on the FST values of SNPs, InDels, and SVs between the Han and Tibetan cohorts. The most Tibetan-specific deletion disrupts an enhancer (red arrow) targeting (yellow line) ATP6V1E2 and RHOQ; it also potentially disrupts a loop (black dot and inverted triangle) boundary. This region shows an additional 121 bp insertion (purple dot) upstream of EPAS1 and SNPs with high LD, possibly reflecting multiple selective events. **d.** Manhattan plot (top), TAD (middle) and haplotype (bottom) near IL1RL1 on chromosome 2 based on the FST values of SNPs, InDels, and SVs between the Han and Tibetan cohorts. A deletion in the exon of IL1RL1 disrupts an enhancer (red arrow) targeting (yellow line) IL1RL1 and IL18RL1; it also potentially disrupts a loop (black dot and inverted triangle in TAD) boundary.

#### SV signals of selection for high-altitude adaptation

Several SNPs are linked to the high-altitude adaptation of Tibetans^5,19–21^; however, the studies on the roles that SVs play in the evolution of Tibetan adaptation to high altitudes are very limited. Therefore, to study the differences in the Han and Tibetan populations, we first filtered the SVs with an F_ST_ >0.1, resulting in 612 SVs. The TibetanSV webserver (https://zhilong.shinyapps.io/tibetan) provides these SVs with associated annotations. Among these SVs, 319 SVs were novel, not being identified in the 1KPG, gnomAD, dbVar and DGV databases, and 457 SVs showed a higher frequency in Tibetan people than in Han people. These SVs are candidate high-altitude adaptation SVs. For example, an insertion was on an intron of *EGLN1* (rank 13, dbsv6981, F_ST_=0.27, 1q42.2), and *EGLN1* is associated with high-altitude, as reported previously^8^. Two intergenic translocations between *MDH1* and *UGP2* (dbsv66039_1 and dbsv66040_1, F_ST_s=0.15, 2p15) were found in 14% of Tibetans and zero Han individuals. *MDH1* was found to be among the top 10 evolutionarily selected regions in rhesus macaques living on a plateau^22^, while *MDG1B*, a paralogue of *MDH1*, has been reported to be a target of selection in Tibetans^4^.

SVs exhibit broad and distal regulation through cis-regulatory circuitry rewiring. Among 612 SVs, 61% SVs (373) overlapped with a promoter, enhancer, silencer, topologically associating domain (TAD) boundary or chromatin loop boundaries. Overall, 71 deletions **(Supplementary Fig. 4b)** and 29 insertions **(Supplementary Fig. 4c)** overlapped with enhancers, 19 SVs overlapped with promoters, 9 SVs overlapped with silencers and 347 SVs were associated with a TAD/loop boundary (see the TibetanSV webserver). For example, one exonic deletion (dbsv59520, F_ST_=0.15, 2q12.1) associated with *IL1RL1* in 45% of Tibetan and 18% of Han individuals resulted in the truncation of one of the protein-coding transcripts of *IL1RL1* (**Fig. 4d**). This deletion, located within a TAD, disrupts a loop and an enhancer in lung tissue, affecting *IL18R1* and *IL1RL* through cis-regulatory circuitry rewiring (**Fig. 4d**). Both *IL1RL1* and *IL18R1* are associated with asthma, a breathing-related lung disease^23^, and many other immune system diseases. The fact that the prevalence rate of asthma in Tibetan children is lower than that in Han children indicates that IL1RL1 could be a target for treating asthma.

We chose 15 SVs with an F_ST_>0.25, including 6 novel SVs, for an in-depth investigation of the relationship between the associated genes and the biological traits of Tibetans (**Fig. 5a**). The biological functions and tissue-specific high expression of the protein-coding genes located near these SVs were visualized in a network (**Fig. 5b**). Some of these genes, including *EPAS1, CRIP2* and *GNPAT*, are highly expressed in artery, lung and heart tissues, while others, such as *UNCX, ADAM19* and *CYFIP2*, are highly expressed in the blood, and many of the genes, such as *TTC7A* and *BRAT1*, are highly expressed in the testis. Importantly, these genes are associated with the response to hypoxia, inflammation, glucose, lipid and energy metabolism, insulin receptor signalling, blood coagulation and keratin filaments in these tissues, indicating their roles in high-altitude adaptation.

**Figure 5.**
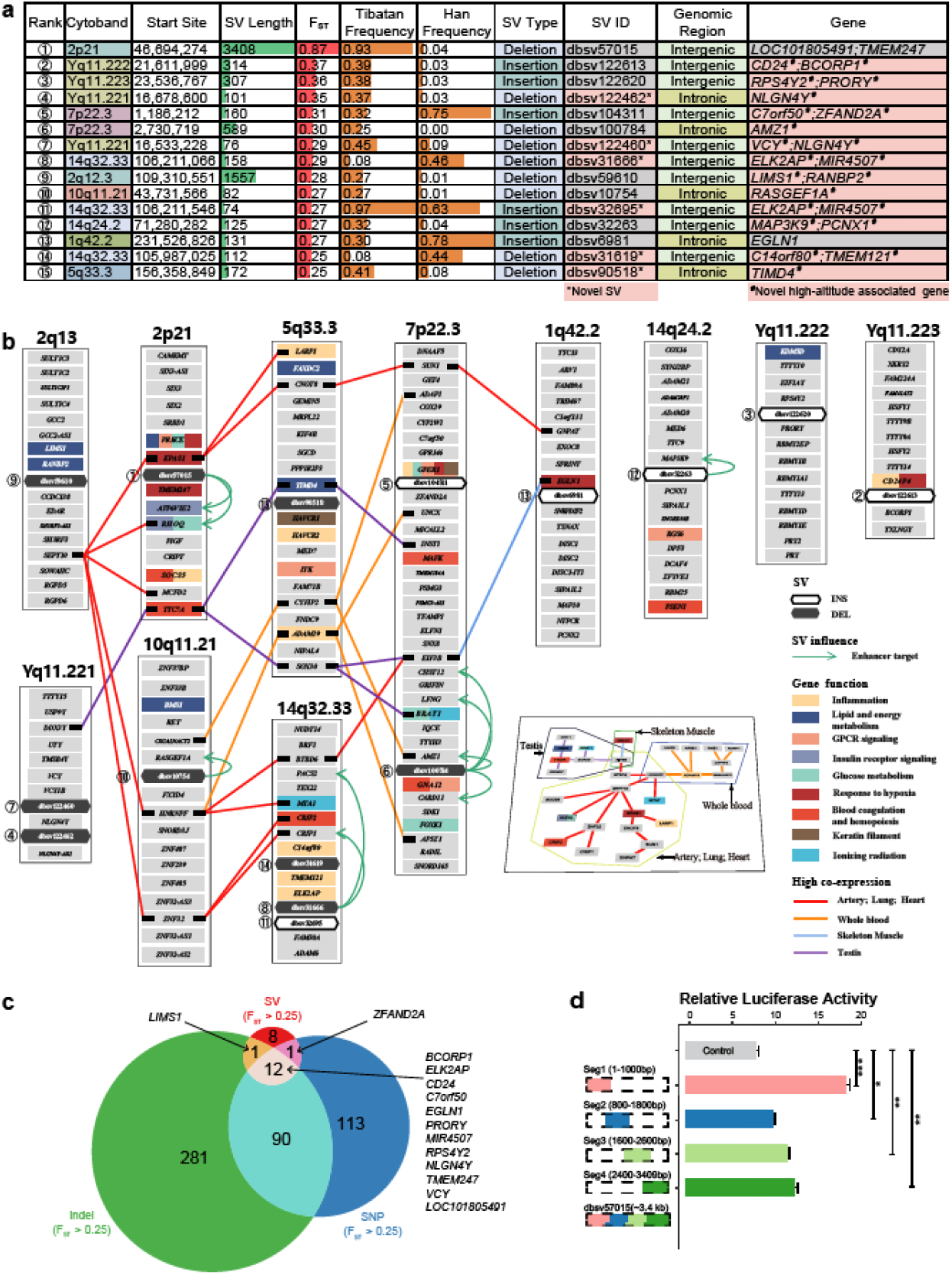
Evolutionary selection of genes for adaptation to high altitude in Tibetans. **a.** Basic description of 15 SVs with an FST>0.25 ordered by FST values. We discovered 6 novel SVs (highlighted in pink in the SV ID column) and 13 novel high-altitude-associated gene groups (highlighted in pink in the Gene column). **b.** The biological functions (coloured rectangles) and tissue-specific high expression (coloured lines) of the protein-coding genes located near the top 15 population-specific SVs (black or white hexagons) are visualized in a network. Most of these genes are related to the response to hypoxia, inflammation, glucose, lipid and energy metabolism, insulin receptor signalling, blood coagulation and keratin filaments in these tissues, indicating their roles in high-altitude adaptation. **c.** Venn diagram of nearby genes related to SNPs, InDels and SVs with an FST>0.25. The high consistency between the associated genes of SNPs, InDels and SVs indicates coincidental natural selection at high altitudes. The non-overlapping genes suggest special functions of these genomic variations. **d.** Fluorescence intensity comparison for the dbsv57015 deletion and control reporter proves that the deleted sequence is an enhancer. Kidney-derived 293T cells were transfected with the pGL3 control vector, Seg1, Seg2, Seg3 or Seg4 (n=5 per group). The pRL-TK plasmid encoding the Renilla luciferase gene was cotransfected into these cells and used as an internal control for transfection efficiency. Both firefly luciferase and Renilla luciferase activities were sequentially measured 48 h after transfection. *P<0.05, **P<0.01, *** P <0.001.

#### EPAS1/TMEM247 SV disrupts a super enhancer

The strongest Tibetan-specific signal was found for the dbsv57015 deletion, located on 2p21 (rank 1, F_ST_=0.87) between two hypoxia-related genes, *EPAS1* and *TMEM247*. This deletion was fully linked (LD=1) with 212 SNPs (78 in *EPAS1*, 19 in *TMEM247*) and 17 InDels in 43 Tibetan samples (**Fig. 4c**), indicating a long haplotype consistent with adaptive selection. Multiple genome-wide association studies (GWASs) have associated these SNPs with high-altitude adaptation, red blood cell counts, body fat distribution, offspring birth weight, and HDL cholesterol (**Supplementary Table 9**), consistent with multiple biological adaptations to high altitude. This region showed an additional insertion (dbsv62730, 2p21, F_ST_=0.20) upstream of *EPAS1* (**Fig. 4c**), possibly reflecting multiple selective events. Notably, both *EPAS1* and *TMEM247* are also positively selected in Tibetan Mastiff dogs^24^ and show distinct association signals^25^, suggesting that they both functionally contribute to adaptation. These results suggest that this SV influences high-altitude adaptation collaboratively with these SNPs and InDels.

We next sought to understand the specific mechanistic role of the dbsv57015 deletion. This deletion overlapped an enhancer predicted by EpiMap^26^ and enhancerDB (vista31415)^27^, indicating a possible gene-regulatory function. We generated 4 truncated segments of the 3.4 kb deletion-containing sequence of dbsv57015 and tested each independently in kidney-derived 293T cells. We found that all 4 sequences showed significantly increased luciferase activity compared to the control (up to 2.3-fold for the first 1 kb segment) (**Fig. 5d**), indicating that the deleted region plays a cis-regulatory role. This deletion also affected the expression of *ATP6V1E2* and *RHOQ*, both of which are targets of the super enhancer (**Fig. 4c and Supplementary Fig. 4b**). These two genes are involved in signalling via GPCR and insulin receptor signalling pathways, which play critical roles in the hypoxia response^28,29^; indeed, both insulin and glucose levels are reduced in Tibetan individuals relative to Han individuals^30^. These findings indicate gene-regulatory roles of dbsv57015 through the deactivation of a gene-regulatory enhancer element with combined effects on multiple genes.

##### The sequence of dbsv57015 affects gene transcription activity via *trans*-regulatory circuitry

A DNA pull-down assay revealed binding proteins including ATP synthase and binding cassette proteins, general transcription factors and prolyl 4-hydroxylase subunit alpha-1 (P4HA1). Surprisingly, P4HA1 expression is induced by HIF-1 under hypoxia, P4HA1 can stabilize HIF-1α, leading to positive feedback regulation and increased expression of HIF-1-induced genes^31^. The binding of P4HA1 indicated a potential role in high-altitude adaptation. Gene Ontology analysis following the DNA pull-down assay showed that the binding proteins of the dbsv57015 sequence were mainly enriched in biological processes related to ribonucleoprotein complex biogenesis and DNA conformation change (**Supplementary Fig. 5 and Supplementary Table 10**). The deletion of the dbsv57015 sequence in Tibetans also rewires the *trans*-regulatory circuitry by disrupting the binding of transcription factors within this genomic region, suggesting a compensatory low demand for oxygen under hypoxia by inhibiting multiple unnecessary molecular and cellular activities. In summary, the most Tibetan-specific deletion was associated with high-altitude adaptation through complex *cis*- and *trans*-regulatory circuitry rewiring.

#### Examples of biological roles for specific SVs

##### Dbsv1O4311 is involved in longevity, sleep quality, and emotion in the Tibetan population

The insertion located between *C7of50* and *ZFAND2A* (rank 5, dbsv104311, F_ST_=0.31, 7p22.3) is accompanied by 5 additional deletions (F_ST_>0.1, TibetanSV webserver), indicating a strong selective signal (**Supplementary Fig. 6a**). These deletions fall within a genetic region that has been associated with insomnia symptoms, longevity, and systolic blood pressure in multiple GWAS (**Supplementary Table 9**). In addition, *ZFAND2A* is associated with the lifespan of *C. elegans*^32^, shows several SNPs with differential allele frequencies in Tibetan individuals (**Fig. 5c**) and has been shown to be selected in Tibetan pigs^24^. *C7orf50* was previously associated with miserableness in a transcriptome-wide association study^33^. Indeed, Tibetan individuals show increased longevity^34^, reduced agrypnia and insomnia symptoms of acute mountain sickness, and reduced dispiritedness relative to non plateau-living individuals, indicating that these SVs may contribute to these adaptive phenotypic differences.

##### Inflammation plays a key role in the survival in cold, hypoxic environments

Three Tibetan-specific SVs were found in the same chromosomal segment (14q32.33), including an intergenic deletion between *ELK2AP* and *MIR4507* (rank 8, dbsv31666, F_ST_=0.27), an intergenic insertion between *ELK2AP* and *MIR4507* (rank 11, dbsv32695, F_ST_=0.27) and a deletion in the intergenic region between *C14orf80* and *TMEM121* (rank 14, dbsv31619, F_ST_=0.25) (**Supplementary Fig. 6b**), indicating a strongly selected region. These SVs are in high LD with multiple SNPs associated with blood protein levels, composite immunoglobulin, and autoimmune traits (**Supplementary Table 9**), indicating potential roles in blood flow and immunoregulation. Indeed, Tibetan individuals show significantly higher total protein levels^30^ associated with inflammation, which is an important response to hypoxic cold environments and can mitigate altitude-related illness^35^.

##### Lipid metabolism supplies additional energy to Tibetans for high-altitude adaptation

A Tibetan-specific intergenic deletion between *LIMS1* and *RANBP2* (rank 9, dbsv59610, F_ST_=0.28, 2q12.3) lies within a region that is genetically associated with triglycerides and LDL cholesterol levels according to GWAS (**Supplementary Table 9**). Indeed, Tibetans exhibit lower triglyceride, cholesterol and LDL levels and higher HDL levels than Han individuals^30^, and they consume more energy generated through lipid metabolism to meet the energetic needs imposed by hypoxia and low temperatures on Han people living on high-altitude plateaus relative to Han individuals living on plains. Indeed, genes related to lipid and fat metabolism have also been associated with high-altitude adaptation in rhesus macaques^22^, indicating convergent metabolic adaptation in humans and other primates. The above deletion is also associated with excessive hairiness, lobe size, and lung function (**Supplementary Table 9**), which are traits consistent with high-altitude adaptation.

##### Several SVs are associated with relieving high-altitude sickness symptoms in Tibetans

One identified Tibetan-specific deletion was located in an intron of *TIMD4* (rank 15, dbsv90518, F_ST_=0.25, 5q33.3), upstream of HAVCR1, a gene associated with headache^36^. This deletion disrupts a predicted enhancer (enh87362) included in EnhancerDB^27^ that is predicted to regulate *TIMD4*. Notably, *HAVCR1* is an important paralogue of *TIMD4*. As headaches and migraines are associated with changes in vascular blood flow to the brain, this deletion may contribute to the observed differences in headache incidence between Han and Tibetan individuals at high altitude^37^. This deletion is also associated with triglycerides, LDL cholesterol and total cholesterol levels (**Supplementary Table 9**), which may provide a mechanistic explanation for these vascular differences or may indicate pleiotropic effects across multiple pathways.

##### Multiple SVs show Tibetan-specific functions in traits such as heel bone mineral density and birth weight

Twelve Tibetan-specific SVs showing an F_ST_>0.1 were associated with heel bone mineral density, including an intergenic deletion between *SLC8A1* and *LINC01913* (rank 21, dbsv58385, F_ST_=0.23, 7p22.3) and an intronic deletion in *CNOT4* (rank 28, dbsv103188, F_ST_=0.22, 7q33) (**Supplementary Table 9**). An intronic deletion in *AMZ1* (rank 6, dbsv100784, F_ST_=0.3, 7p22.3) disrupts an enhancer whose targets include *AMZ1*, which is associated with heel bone mineral density according to the GWAS Catalog, possibly contributing to the increase in tP1NP procollagen^30^ and lower prevalence of osteoporosis observed in Tibetans relative to the Han population.

Ten identified SVs with an F_ST_>0.1 are associated with birth weight (**Supplementary Table 9**). Generally, high-altitude reduces infant birth weight as a result of intrauterine growth restriction; however, the birth weight of Tibetans is higher than that of Han individuals living at high altitudes^38^.

##### Other SVs of unclear function that are worth exploring

Among the 15 SVs with an F_ST_>0.25, one intronic SV and one intergenic SV were associated with *NLGN4Y* (**Supplementary Fig. 6c**), which is related to learning, vocalization behaviour, presynapse assembly, and autism. Two of the intergenic SVs were associated with *ELK2AP, MIR4507* or *IGHG1* and *IGHG3* according to GENCODE annotation (**Supplementary Fig. 6b**). *RPS4Y2* (**Supplementary Fig. 6d**) and *RANBP2* (**Supplementary Fig. 6e**) are involved in influenza viral RNA transcription and replication. Additionally, *RANBP2* is related to the regulation of glucokinase and hexokinase activity. *CD24* (**Supplementary Fig. 6f**), *MAPK3K9* and *RASGEF1A* are involved in the MAPK cascade.

##### Gene variations, including SNPs, InDels, and SVs, regulate genes both collectively and independently

As we found that several peaks in the Manhattan plot of SNPs, InDels, and SVs were highly consistent (**Fig. 4a**), the collection of SNPs, InDels and SVs with an F_ST_ >0.25 provides a set of perfect candidate genes that may be involved in the high-altitude adaptation of Tibetans. We obtained overlapping genes between the annotated genes related to SNPs and InDels with an F_ST_ > 0.25, and 12 of the 22 genes appeared in all 3 gene sets (**Fig. 5b**). The overlap between the three call sets verified that most of our identified SVs are associated with high-altitude adaptation. A total of 382 genes showing an F_ST_ greater than 0.1 were identified among the genes associated with SVs, SNPs and InDels. Although there were different genes in each set, the overlapping genes between them indicated that SVs, InDels and SNPs function collectively to support high-altitude adaptation. However, different types of gene variations were also shown to exhibit specific functions when considering non-overlapping genes. This suggests both cooperative and independent contributions of SVs, SNPs, and InDels in high-altitude adaptation. More generally, gene variations, consisting of SNPs, InDels, and SVs, function both collectively and independently to regulate genes in human biology.

##### Denisovan-introgressed SVs were not selected during the evolution of the Tibetan genome

We also identified several ancient SVs that existed in apes or ancient hominins (**Supplementary Fig. 7**). For example, the *EGLN1*-related insertion (rank 13, dbsv6981, F_ST_=0.27, 1q42.2) was identified from apes; a duplication in the exonic *HLA-DRB5* (rank37, dbsv99370, F_ST_=0.22, 6p21.32) was identified in Denisovans and did not exist in apes, Africans, or Neanderthals. Notably, no SVs with high F_ST_ values were discovered in ancient hominins, although a 5-SNP motif in *EPAS1* introgressed from Denisovans contributed to the high-altitude adaptation of Tibetans.

#### Functional significance of SVs in evolutionary adaptation

The high-altitude environment shapes the fat metabolism, steroid hormone production, heart functions, and brain development of Tibetans to allow them survive on the plateau. Beyond these individual examples, we searched for systematic genome-wide enrichment of our Tibetan (**Supplementary Fig. 8a**) and Han (**Supplementary Fig. 8b**) SVs in specific pathways through KEGG^39^ analysis, excluding singleton SVs. We found that Tibetan-specific SVs were enriched in multiple key metabolic pathways, consistent with the adaptive advantages of Tibetans in cold and hypoxic environments, including those related to steroid hormone biosynthesis and fat digestion and absorption, which can be helpful for producing sufficient body heat and energy in cold environments. We also observed enrichment related to vascular function, pulmonary function, blood pressure, and vascular smooth muscle contraction, which can play important roles in adaptation to hypoxic conditions. We observed enrichment in alpha-linolenic acid metabolic pathways, which are known to reduce the risk of coronary heart disease and improve cardiovascular health. We found strong enrichment of the herpes simplex virus 1 infection pathway, consistent with the fact that the prevalence and incidence of herpes simplex virus 1 infections are relatively higher in the Western Pacific^40^, likely reflecting a historic prevalence of viral infections. We also found notable enrichment of several Gene Ontology^41^ terms. These terms were related to synapse assembly and organization, cognition, and learning or memory processes (enriched in Tibetan-specific SVs with an F_ST_ >0.1) and several immune processes, including T cell activation (enriched in Han-specific SVs with an F_ST_ >0.1 (**Supplementary Fig. 8c**). These genes also showed enrichment related to specific Gene Ontology terms related to cellular components, including synaptic localization (**Supplementary Fig. 8d**), consistent with observed differences in cognitive pathways. These broad biological enrichment results indicate that high-altitude adaptation involves multiple biological pathways related to metabolism, vasculature, circulation, and cognition allowing survival in cold, hypoxic environments.

## Discussion

Our study provides an important high-resolution view of the high-altitude adaptation of Tibetans based on the long-read sequencing of 320 Han and Tibetan genomes, revealing the complex SV landscape of Han and Tibetan populations, and we further obtained important insights for understanding the evolutionary adaptation of the Tibetan population through a systematic study of the genomic SV landscape. We provide 122,876 high-quality Han and Tibetan SVs, dramatically expanding the known landscape of genetic variation and the corresponding resources available for East Asian populations. We revealed many candidate SVs and genes for high-altitude adaptation, revealing diverse biological adaptations consistent with the observed physiological differences in the Tibetan population. The most Tibetan-specific SV has a broad impact on gene regulation via complex *cis*- and *trans*-regulatory circuits. Different types of genomic variations, consisting of SNPs, InDels, and SVs, function both in combination and in parallel.

The understanding of the human genome is changing dramatically with continuing technological developments. The NGS of large cohorts has revealed the important roles of genomic variations in the evolutionary adaptation of human populations. Our long-read ONT-based study provides the first large SV reference panel based on a cohort of 320 Han and Tibetan individuals. Quality evaluation of our SV call sets through comparisons between our SV call set and those of the 1KGP and gnomAD, comparisons between ONT and CCS SV call sets from the same sample, and qPCR validation confirmed the high quality of this call set. It fills a gap in lower EAS-based genetic resources for community studies, as reported previously. Our study further confirmed the high enrichment of SVs in genomic repeat regions. We also found hotspots of SVs in the genome and showed an evolutionary preference for repeat regions associated with intronic SVs. All these results reveal a prospective landscape of high genetic diversity and complexity for human genomic variations and evolution.

We systematically assessed the SV landscape of Han and Tibetan populations. The Han and Tibetan populations could be separated by PCA and admixture analysis based only on the SV call set. This situation has previously been demonstrated through SNP analysis^4^. This finding demonstrated that SVs can be considered functionally equal to SNPs to some extent. SVs and their impacts on human health and diseases are worthy of broad, in-depth studies. We compared the SVs between Han and Tibetan populations and identified SVs with high F_ST_ values, which are probably related to high-altitude adaptation. Several of these SVs are related to known hypoxia-associated genes, while most of them were have not been previously identified in high-altitude environments, and six of the SVs were novel. These findings proved the value of the ONT resequencing of the Han and Tibetan populations. Our results provide a great resource for the identification of candidate genes for high-altitude adaptation.

The selection observed among SNPs, InDels, and SVs was highly consistent between the Tibetan and Han populations. By comparing genes related to SNPs, InDels, and SVs with high F_ST_ values between the Han and Tibetan populations, we found that some genes were simultaneously associated with the three types of gene variation. This provides a strong signal that these genes are involved in high-altitude adaptation. There were also some genes that were associated with only one or two types of gene variation. Different types of gene variation function both in combination and in parallel to achieve perfect adaptation to the plateau environment.

We found that SVs show wide and distal regulation through cis- and trans-regulatory circuitry rewiring. A majority of the SVs with an F_ST_ >0.1 overlapped with *cis*-regulatory elements. An *IL1RL1*-associated exonic deletion disrupts an exon of *IL1RL1* and the enhancer, suggesting the disruption of this gene and *cis*-regulatory circuitry rewiring. Multiple SVs, including the most Tibetan-specific SVs and an *EGLN1*-associated SV, disrupt a TAD or loop boundary, affecting the cis-regulatory circuitry. We used an enhancer reporter assay and a DNA pull-down assay to show that the strongest Tibetan-specific SV (F_ST_=0.87) is associated with a complex *cis*- and *trans*-regulatory circuit, resulting in the deletion of a super enhancer bound by several key transcription factors and targeting multiple nearby genes through proximal and distal interactions, illustrating the role of non-exonic SVs in gene-regulatory circuitry rewiring.

We provide several dramatic examples of adaptation related to the hypoxia response, red blood cell count, blood pressure, body fat distribution, birth weight, bone mineral density, energy and lipid metabolism, insomnia, agrypnia, longevity, mountain sickness, immunoregulation, inflammation, lung function, brain vascular blood flow, and headache. These examples illustrate the broad set of biological processes involved in high-altitude adaptation, the biological relevance of our findings, and the power of our integrative genomics approach for revealing the biological processes involved in adaptive events.

Overall, our study demonstrates the power of single-molecule long-read sequencing, provides an important greatly expanded comprehensive reference for global SVs, reveals many dramatic biological examples of human adaptation, provides important biological targets for combatting hypoxia, illustrates complex gene-regulatory circuitry rewiring mediated by SVs, and provides a wealth of biological insights into human biology and recent human adaptation.

## Supplementary figure legends

**Supplementary Figure 1. Han & Tibetan SV call set.** (a) The number distribution of each type of SVs in Han and Tibetan samples. (b) The cumulative percentage of the number of SVs excluding singletons within the Tibetan cohort. (c) The cumulative percentage of the numzber of SVs excluding singletons within the Han cohort. (d) The cumulative percentage of the number of SVs excluding singletons within the Han and Tibetan cohort. (e) Venn diagram between the Han and Tibetan SV call set and the SVs identified based on 15 publicly available genomes generated by long-read sequencing. (f) Venn diagrams of the overlap among the Han and Tibetan, 1KGP, gnomAD and 15 genome SV call sets.

**Supplementary Figure 2. Characteristics of SV distribution and composition.** (a) Frequencies of different types of SVs with lengths shorter than 1 kb. The length of Alu is approximately 300 bp. (b) Frequencies of different types of SVs with lengths longer than 1 kb. The length of LINEs is approximately 6 kb. (c) The numbers of 5 types of SVs in different gene elements. (d) The proportions of SINE- and LINE-mediated SVs and exonic SVs in different types of SVs. (e) The frequencies of SVs and the proportions of all SVs. (f) Overall, 58 SVs are associated with C7orf50. (g) The ratio between repeat-associated SVs and non-repeat associated SVs among singleton, all and shared SVs. (h) The percentages of different gene elements related to singleton, all and shared SVs in the Han and Tibetan populations. (i) The percentages of 5 types of SVs related to singleton, all and shared SVs in the Han and Tibetan populations.

**Supplementary Figure 3. Population genetics of Han and Tibetan populations.** (a) Overlap between the SVs in Han and Tibetan populations, showing that they share a majority of SVs. (b) PCA of the SV call set of the Tibetan (green) and Han (red) cohorts indicates clear separation between the two groups.

**Supplementary Figure 4. Chromosome landscape of Han-Tibetan population and related enhancers.** (a) Manhattan plot for each chromosome based on the F_ST_ values (y-axis) of SVs (orange-red boxes), SNPs (blue green dots) and InDels (dark blue triangles) between the Han and Tibetan cohorts. (b) The SVIDs of deletions and genes targeted by enhancers (y-axis) are connected (red) with different enhancers in different tissues (x-axis). (c) The SVIDs of insertions and genes targeted by enhancers (y-axis) are connected (red) with different enhancers in different tissues (x-axis).

**Supplementary Figure 5. DNA pull-down results for the dbsv57015 sequence.** (a) The sequence of dbsv57015 consists of T1 and T2. (b) Venn diagram of pulled-down proteins in the 293T cell line. (c) Gene Ontology (GO) biological process enrichment of the proteins in the control and pull-down groups in the 293T cell line. (d) Venn diagram of pulled-down proteins in the U266 cell line. (e) GO biological process enrichment of the proteins in the control and pull-down groups in the U266 cell line.

**Supplementary Figure 6. Manhattan plot of several population-specific genomic regions.** (a) Manhattan plot of the region near the *ZFAND2A* gene based on the F_ST_ values of SVs, SNPs and InDels between the Han and Tibetan cohorts. (b) Manhattan plot near gene *ELK2AP* based on the F_ST_ of SV, SNP and InDel between the Han and Tibetan cohorts. (c) Manhattan plot of the region near the *NLGN4Y* gene based on the F_ST_ values of SVs, SNPs and InDels between the Han and Tibetan cohorts. (d) Manhattan plot of the region near the *PRS4Y2* gene based on the F_ST_ values of SVs, SNPs and InDels between the Han and Tibetan cohorts. (e) Manhattan plot of the region near the *RANBP2* gene based on the F_ST_ values of SVs, SNPs and InDels between the Han and Tibetan cohorts. (f) Manhattan plot of the region near the *CD24* gene based on the F_ST_ values of SVs, SNPs and InDels between the Han and Tibetan cohorts.

**Supplementary Figure 7. Possible evolutionary scenarios of SVs.** The numbers of SVs are shown under each scenario. In addition, a majority of SVs existed only in the Tibetan and Han populations.

**Supplementary Figure 8. Enrichment analysis of Han/Tibetan-specific SVs.** (a) Top 10 KEGG pathways of Tibetan-specific SVs, excluding singleton SVs. (b) Top 10 KEGG pathways of Han-specific SVs, excluding singleton SVs. (c) Gene Ontology (GO) biological process enrichment terms for Han-specific or Tibetan-specific SVs with an F_ST_>0.1. (d) GO cellular component enrichment terms of Han-specific or Tibetan-specific SVs with an F_ST_>0.1.

## Supplementary Table legends

**Supplementary Table 1.** (a) Samples and ONT sequencing statistical information. (b) Manual curation of 298 SVs across all samples. (c) PCR validation of 4 SVs in 57 samples. (d) Statistics of SMRT-CCS sequencing data. (e) The number and frequency of different gene elements and types of SVs among singleton, shared and all SVs. (f) Samples and NGS statistical information.

**Supplementary Table 2.** Statistics of ONT and SMRT-CCS sequencing data.

**Supplementary Table 3.** Orthogonal validation of ONT-SVs against SMRT-CCS-SVs from the same sample.

**Supplementary Table 4.** AF distribution of different types of novel SVs in the Tibetan-Han population.

**Supplementary Table 5.** SV comparison between ZF1 and our Tibetan population data.

**Supplementary Table 6.** Mean SV statistics for each sample of different AFs in the Tibetan-Han population.

**Supplementary Table 7.** SV distribution in different genomic regions in the Tibetan-Han population.

**Supplementary Table 8.** SINE- and LINE-associated SVs in various genomic functional regions.

**Supplementary Table 9.** SV-SNP-GWAS-phenotype analysis results.

**Supplementary Table 10.** Proteins identified and their annotation in DNA pull-down assays of the sequence of the most Tibetan-specific SVs.

## Method details

### Key resources table

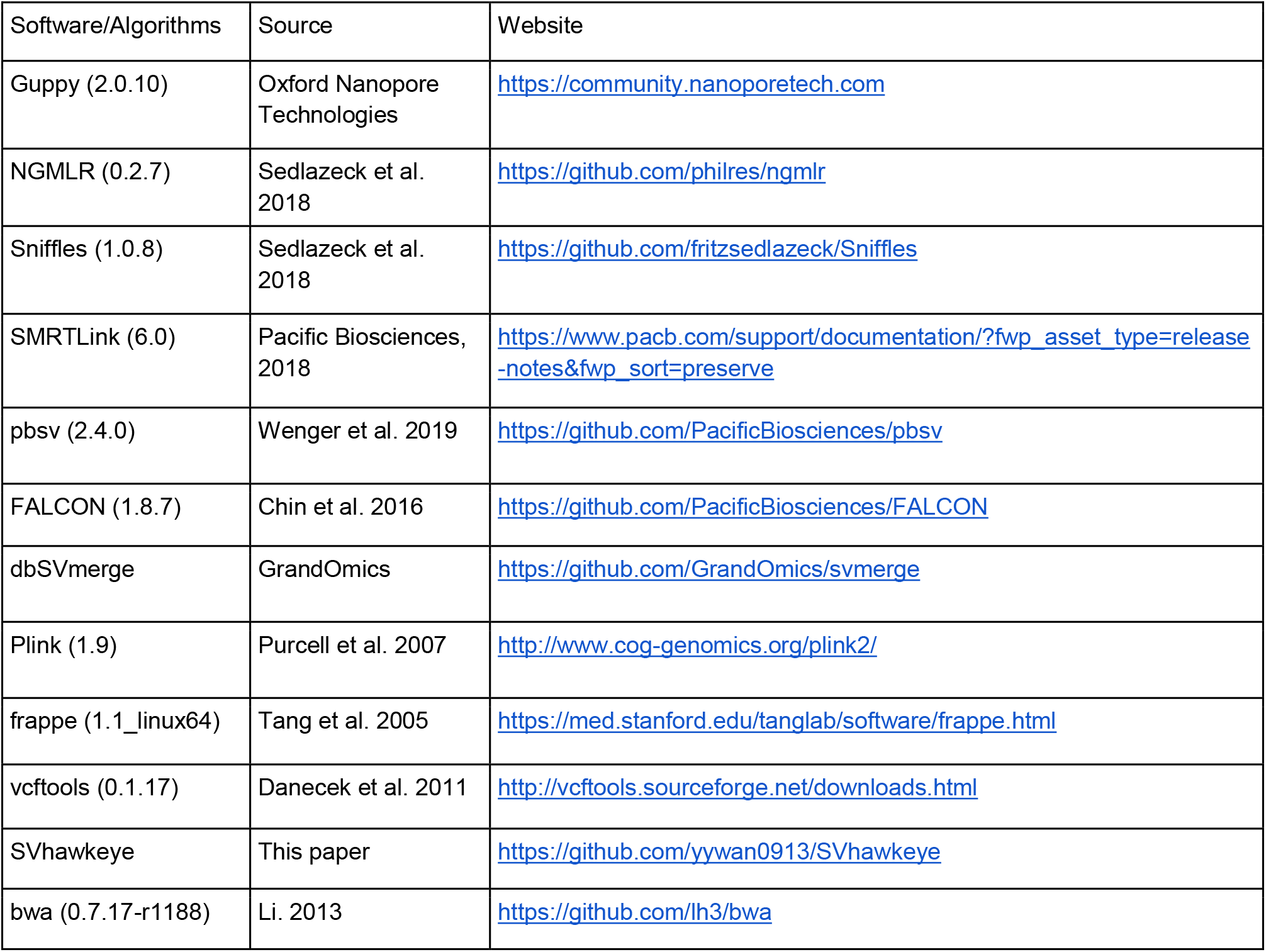
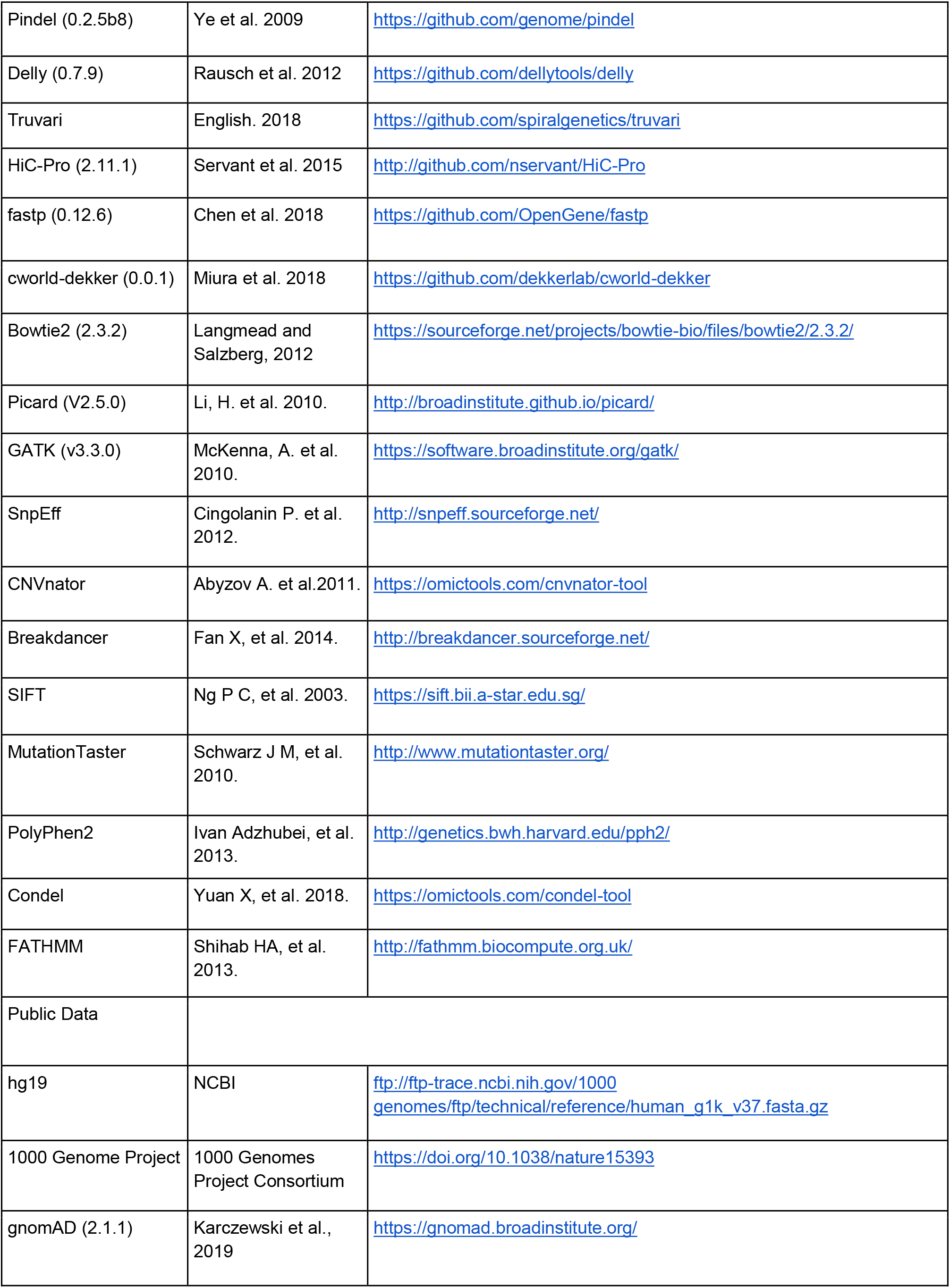
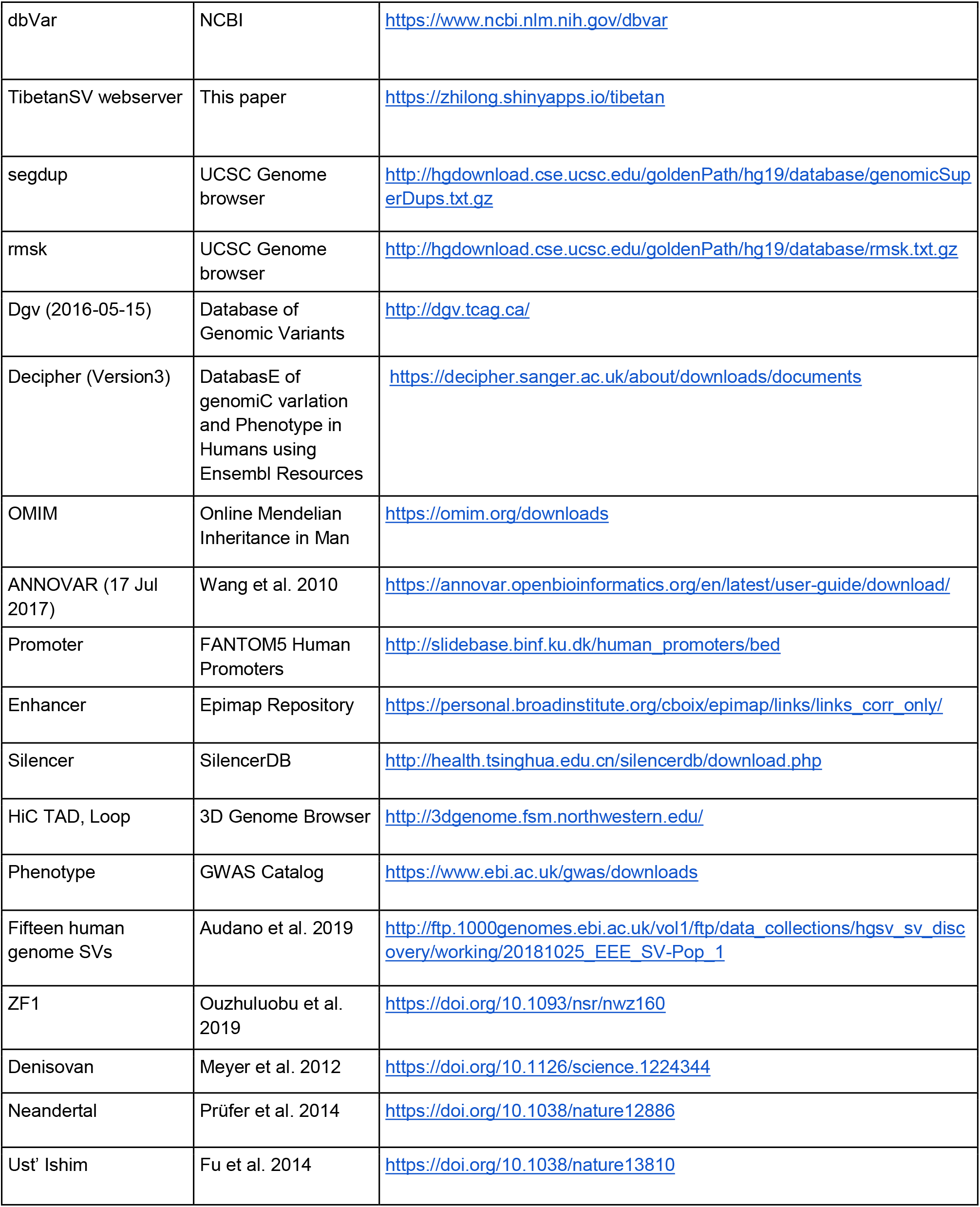

#### Samples and third-generation sequencing

To ensure that we covered the majority of SVs that were shared within or among the Han or Tibetan populations, we sequenced 201 Han and 119 Tibetan genomes based on ONT sequencing. All the samples were selected randomly (**Table S1**). We oversampled Han genomes by sequencing more individual Han genomes than Tibetan genomes because we anticipated finding more diverse SVs in the Han population, which shows high admixture relative to the Tibetan population.

Genomic DNA was prepared from each of the 320 samples by sodium dodecyl sulphate (SDS)-based methods. Then, nanopore libraries were constructed according to the manufacturer’s instructions for the Ligation Sequencing Kit 1D (SQK-LSK109) and sequenced on R9.4 flow cells using a PromethION sequencer (ONT, UK) at the Genome Center of Grandomics (Beijing, China). Base calling was subsequently performed from fast5 files using Guppy (v2.0.10) software to generate the FASTQ files.

To compare the results of SV calling with those of the other long-read sequencing platform, one of the Tibetan samples, AL-2-33, was randomly selected for sequencing on the PacBio Sequel system. The genomic DNA was sheared, and size selection of 10-15 kb fragments was performed by using BluePippin (Sage Science, USA). SMRTbell libraries were constructed using the SMRTbell Template Prep Kit v.1.0 (100-259-100, PacBio) and then sequenced using V3.0 chemistry on the PacBio Sequel system. CCS reads were generated using SMRTLink (v6.0) software from PacBio.

#### SV calling

To produce high-quality reference SV sets for the Han and Tibetan populations, we constructed a stringent analysis workflow. The complete sequencing and analysis workflow consisted of 1) an average sequencing depth of 20.85±7.54X with a long read length (average N50 length=22.71±4.04 kb) (**Table S1A**); 2) SV calling by using Sniffles, which has been proven to be the most effective computational tool for identifying SVs from nanopore sequencing data thus far^13^; 3) the orthogonal validation of ONT-SVs against SMRT-CCS-SVs using 453.79 Gb of total polymerase bases, which produces CCS data with a 15.4X depth (**Table S1D**); 4) the manual curation of 95,360 SVs from 298 candidate regions (**Table S1B**) that were originally mapped by the computational pipeline; 5) the selection of 228 SVs (**Table S1C**) for validation using PCR and Sanger sequencing; and 6) the cross-comparison of our SVs with those in various human genome variant databases and those reported in recently published work. The FASTQ files were aligned to the hg19 human reference genome available from NCBI, and a BAM format alignment file was produced using NGMLR v0.2.7^13^ with default parameters, which is designed for long-read mapping. Variants were called using Sniffles v1.0.8^13^ with the parameter setting -q 0 -l 50 --report_BND, and VCF format files were generated, which contained all SV information. Then, raw SVs were filtered and merged to obtain high-quality SVs. The two filtering criteria were that the AF needed to be greater than 0.3 and that the sequencing depth of the variant had to be greater than the larger number between 0.3-fold and 2-fold the average sequencing depth and less than the twice the average sequencing depth. High-quality SVs were used for the subsequent analysis. The SVs derived from PacBio and ONT sequencing platforms were compared according to the following criterion: 50 bp ≤ SV length ≤50,000 bp, using Truvari.

#### SV annotation and distributions

High-quality SVs with upstream and downstream genes were annotated in the segdup, rmsk, dgv, 1000 Genome Project, gnomAD, Decipher, OMIM, and ANNOVAR databases. We grouped the SVs into 500 kb bins to count the number of various types of SVs (insertions, duplications, deletions, inversions, and translocations). The cytoband file was downloaded from the UCSC website, and chromosome banding was drawn for different regions. The numbers of SVs indicated in different colours were determined using the R language 3.4.1^42^

#### SV merging

DbSVmerge was used to obtain the nonredundant SV set for all high-quality SVs from 320 samples. The merging strategy was as follows: the distance of the variant coordinates between any two SVs must be less than 1 kb; for deletion-, duplication- and inversion-type SVs, at least 40% of the region should overlap; and for insertion SVs, the difference in length between insertions should be less than twice the length of both insertions. The numbers and lengths of SVs of different types (insertions, duplications, deletions, inversions, and translocations) were counted.

#### SVs comparison among published datasets

We compared our SV calls to several published datasets, including the 1KGP dataset^43^, gnomAD 2.1.1, SVs from fifteen human genome sequences obtained on PacBio platforms [8], and Tibetan ZF1 SVs^16^. The same merging strategy was applied using dbSVmerge. The SV comparison was conducted with Truvari according to the following criterion: 50≤ SV length ≤50,000.

#### SV landscape in Tibetan and Han Chinese populations

We constructed nonredundant sets of 93,154 SVs from 119 samples from the Tibetan population and 109,438 SVs from 201 samples from the Han Chinese population. In this study, SV frequency (also called allele frequency (AF)) was defined as the proportion of the sample size with one SV in the population (calculated excluding singletons). To determine whether the sample size was large enough for SV population analysis, we drew curves of SV counts among samples as every sample was added to the population.

We divided AFs into 5 levels: 0~0.1, 0.1~0.4, 0.4~1, and 1. SV numbers were counted according to the 5 levels and the singletons in each individual. Then, the diversity of the SVs in repeat and nonrepeat regions was quantified for different types of SVs.

#### Genome evolution analysis using SVs

Each SV was given a value of 1 if someone had it and 0 if not, resulting in an N×M matrix, where N stands for the number of samples (320) and M represents the total number of all SVs. All PCAs, evolutionary trees, and population structure analyses were based on the N×M matrix. PCA was carried out and the principal component values were calculated with the R 3.4.1 prcomp function, ensuring that fewer principle components were reserved than the number of samples. Hierarchical clustering was included for plotting the evolutionary tree using the R 3.4.1 hcluster function. Population genetic structure analysis can reveal the time span of the development of subgroups by dividing a large population into several subgroups. Plink^44^ software in two modes, pep and map, was utilized to obtain structural information statistics for each individual, which were analysed with Frappe software^45^.

#### Short-read sequencing, SNP and InDel calling

Short-read sequencing of 148 samples (75 Tibetan and 73 Han, including 113 DNA samples (previously used for ONT sequencing) and 35 blood samples) was performed after a series of sample and library processing steps. A Qubit Fluorometer was used to evaluate the concentration of DNA, and agarose gel electrophoresis was used to examine sample integrity and purity. Fragmented DNA was obtained through Covaris preparation and subjected to selection at an average size of 200-400 bp using an Agencourt AMPure XP-Medium kit. The PCR-amplified products were recovered with the AxyPrep Mag PCR clean up kit.

We performed the paired-end sequencing of the 148 samples with an average output of 137 Gb raw bases. Each sample showed a read depth >31.9X. To reduce sequencing noise, we removed reads containing a) 10% or more ‘N’ bases, b) 50% or more low-quality bases, or c) sequencing adapters. After data filtering, we applied Burrows-Wheeler Aligner (BWA v0.7.12) to map the clean reads against the hg19 human reference genome. We sorted the mapping results and marked duplicate reads in BAM files using Picard tools (v1.118). We performed base quality score recalibration (BQSR) and local realignment around InDels to obtain a more accurate base quality and therefore improve the accuracy of the variant calls. We detected SNPs and small InDels using HaplotypeCaller from the Genome Analysis Toolkit (GATK, v3.3.0). We further applied variant quality score recalibration (VQSR), a variant filtering tool based on the machine learning method, to obtain reliable variant calls with high confidence.

#### Analysis of adaptive evolution using SVs

F_ST_ measures population differentiation due to genetic structure using the SV frequency, calculated as the difference between total heterozygosity and average population heterozygosity divided by total heterozygosity. The heterozygosity frequency of SVs between populations was calculated by Weir and Cockerham estimators using the statistical method of VCFtools^46^.

Tibetan-specific SVs were selected on the basis of satisfying three criteria: the population frequency in Tibetans was not less than 0.2, the population frequency in Tibetans was twice as high as that in Han individuals, and the F_ST_ of the SV was larger than 0.1. Han-specific SVs were obtained using a similar strategy: the population frequency in the Han population should be not less than 0.2, the population frequency in the Han population should be twice that in the Tibetan population, and the F_ST_ of SV should be larger than 0.1. Some population-specific SVs were examined manually with IGV^47^.

Raw data, either FASTQ or BAM files, for Denisovan^48^, Altai Neandertal^49^, Vindija Neandertal^50^ and Ust’ Ishim genomes^51^ were downloaded from the European Nucleotide Archive. We aligned the archaic short reads to the hg19 human reference genome using BWA-MEM^52^. For each archaic hominin genome, we applied Pindel^53^ and Delly^54^ to detect SVs and merged the results from the two SV callers using SURVIVOR^55^.

The SV call set from great ape genomes was downloaded from the Database of Genomic Variants archive (accession number estd235). We converted the coordinates of SVs from reference genome hg38 to hg19 with the modified open-source tool CrossMap^56^. We applied dbSVmerge to merge all of these SV call sets with our own SV call set to determine whether an SV was shared between two genomes or groups/species.

#### LD and GWAS computation

To explore the possible connections between different variations (SV and SNP, InDel), the VCF results of 113 samples that were matched with ONT sequencing samples and the VCF results of 320 SVs were used. The PLINK program was applied to compute the linkage disequilibrium value (LD, R^2^). R^2^ values were computed between SVs and SNPs (InDels) within a window of 1 M bp. The cut-off was set as 0.2. To annotate the functions of SVs according to SNPs found to be associated with various phenotypes through GWASs, we made use of the NHGRI Catalog of published GWASs, which includes 159,203 SNPs linked to a multitude of phenotypes. Finally, 1,632 SVs were shown to be in strong LD (R^2^ ≥ 0.8) with a GWAS SNP found in Han populations, and 1455 SVs were in strong LD (R^2^ ≥ 0.8) with a GWAS SNP found in Tibetan populations.

#### The associations between SV, promoter, silencer, enhancers and HiC data

We downloaded gene-enhancer link data from 833 samples in the Epimap Repository^26^. To provide some tolerance, we extended the breakpoints of 612 SVs (F_ST_ > 0.1) by +-100 bp. We intersected these SV regions with enhancer regions in each sample. Overall, 100 SV regions showing overlap with at least one enhancer in any sample were visualized in a heatmap.

We analysed the associations between SVs and promoters, silencers, and LADs/TADs/loops in a similar way. The promoter data were downloaded from FANTOM5 Human Promoters, and the silencer data were downloaded from SilencerDB. LAD domain data were downloaded from Roadmap, and TAD/Loop data were obtained from the 3D Genome Browser.

#### Pathway and co-expression annotation

We selected the 15 SVs with an F_ST_>0.25 and identified the ten upstream and downstream genes of these SVs (excluding noncoding genes) using the GENCODE v29 GRCh37 gff file. According to the TPM data of these genes in the GTEx database (V8 release), we chose the top ten genes that were highly expressed in the heart, artery, lung, testis, and whole blood as the genes showing high coexpression in the tissue. To illustrate the relationships between these genes and high-altitude adaptation, we chose the GWAS Catalog, SuperPath (https://pathcards.genecards.org/), Gene Ontology, and KEGG databases to annotate the functions of these genes. The clusterProfiler^57^ package was used to enrich pathways.

#### Statistical analysis

All statistical analyses were performed using the R package (v3.4.1, http://www.r-project.org/).

#### Dual-luciferase reporter gene assay to validate the function of the dbsv57015 sequence

We employed a dual-luciferase reporter gene assay to investigate whether the 3.4 kb dbsv57015 deletion plays a role as a cis-element. For this reason, the four truncated sequences representing different base pair truncations (Seg1: 1-1000 bp; Seg2: 1600-2400 bp; Seg3: 2600-3409 bp; Seg4: 800-1800 bp) were cloned into the pGL3-control plasmid (Promega, Madison, WI, USA), upstream of the SV40 promoter and the firefly luciferase reporter gene. A 200-base pair overlap sequence between two truncated sequences was designed to avoid the disruption of an enhancer, as the core length of an enhancer is approximately 100-200 bp. These truncated TED-luciferase plasmids were transfected into 293T cells. Additionally, the pRL-TK vector (Promega, Madison, WI, USA), encoding Renilla luciferase, was cotransfected in combination with dbsv57015-luciferase reporters as an internal control. Both firefly luciferase and Renilla luciferase activities were sequentially measured 48 h after transfection. Firefly luciferase activity was normalized to Renilla luciferase for each sample.

#### DNA pull-down assay to identify the trans-acting regulators of the dbsv57015 sequence

The DNA sequence of dbsv57015 was separated into two parts: T1 (1-1725 bp) and T2 (1676-3409 bp). T1 and T2 DNA probes were affixed to streptavidin magnetic beads (Beaver Biosciences Inc. China) and then incubated with 293T and U266B1 cell lysates at 4°C overnight. We washed the beads on a magnetic rack with buffers containing nonspecific DNA and a low salt concentration (50 mmol/L Tris-HCl, pH 7.6), which removed nonadhering and low-specificity DNA-binding proteins. Then, we washed the beads with higher salt concentrations (100 mmol/L Tris-HCl, pH 8.5) to elute specific DNA-binding proteins, which were used to perform sodium dodecyl sulphate polyacrylamide gel electrophoresis. Silver-stained protein bands corresponding to blank beads (control) and T1 and T2 DNA sequences were used for mass spectrometry analysis.

#### Data availability

The SV dataset supporting the conclusions of this article is available in the Genome Sequence Archive repository as accession PRJCA004371(GVM000124, GVM000125) upon acceptance. The data utilization has an ethical filing number from the Human Genetic Resource Administration of China (2020BAT0405). The study was approved by the Medical Ethical Committee of Chinese PLA General Hospital (Beijing, China, S2018-298-02).

## Acknowledgements

The work was partly supported by the National International Science & Technology Cooperation Special Program [2013DFA31170] and the Science and Technology program of Beijing [Z151100003915075]. We thank the volunteers who donated samples for this meaningful study. We thank the strong support from Prof. Tian Yaping, Wang Weidong. We also thank all the participants who contributed to this study. We thank all the professional comments and suggestions from Ming Ni at Beijing Institute of Radiation Medicine, Qiang Qiu at Northwestern Polytechnical University, Dongdong Wu at Kunming Institute of Zoology, Rasmus Nielsen at UC Berkeley, Asan at BGI-Shenzhen.

## Author contributions

Conceptualization, K.He; Methodology, K.He, J.Shi., Zh.Jia, J.Sun and F.Liang; Study participants recruitment, X.Zhao, J.Shi., Zh.Jia and Q.Jia; Sample collection and preparation: X.Zhao, J.Shi., Zh.Jia, Q.Jia, K.Yu, Sh.Wu, S.Cui, Q.Zhong and J.Wu; Data generation and analysis, Zh.Jia, J.Shi., J.Sun, F.Liang, Ch.Zhao, Depeng Wang, Y.Xiao, Y.Liu and Zh.Wu; Annotation and Functional Links: K.He, M.P, J.Shi., Zh.Jia, J.Sun, Ch.Zhao, X.Song and Q.Chen; Experimental Validation: K.He, M.P, X.Wang and Zh.Jia; Writing and revision: J.Shi, Zh.Jia, M.P, F.Liang, M.K., X.Bo and Zh.Wu; Ethics application and data resource: K.He, J.Shi, K.Yu and X.Zhao; Supervision, K.He.

## References

1. Spielmann, M., Lupiáñez, D. G. & Mundlos, S. Structural variation in the 3D genome. Nat. Rev. Genet. 19, 453–467 (2018).

2. Laugsch, M. et al. Modeling the Pathological Long-Range Regulatory Effects of Human Structural Variation with Patient-Specific hiPSCs. Cell Stem Cell 24, 736–752.e12 (2019).

3. Perry, G. H. et al. Diet and the evolution of human amylase gene copy number variation. Nat. Genet. 39, 1256–1260 (2007).

4. Yi, X. et al. Sequencing of 50 human exomes reveals adaptation to high altitude. Science 329, 75–78 (2010).

5. Lorenzo, F. R. et al. A genetic mechanism for Tibetan high-altitude adaptation. Nat. Genet. 46, 951–956 (2014).

6. Huerta-Sánchez, E. et al. Altitude adaptation in Tibetans caused by introgression of Denisovan-like DNA. Nature 512, 194–197 (2014).

7. Chen, F. et al. A late Middle Pleistocene Denisovan mandible from the Tibetan Plateau. Nature 569, 409–412 (2019).

8. Simonson, T. S. et al. Genetic evidence for high-altitude adaptation in Tibet. Science 329, 72–75 (2010).

9. Genetics for all. Nat. Genet. 51, 579 (2019).

10. Audano, P. A. et al. Characterizing the Major Structural Variant Alleles of the Human Genome. Cell 176, 663–675.e19 (2019).

11. Rishishwar, L., Tellez Villa, C. E. & Jordan, I. K. Transposable element polymorphisms recapitulate human evolution. Mob. DNA 6, 21 (2015).

12. Lin, Y.-L. & Gokcumen, O. Fine-Scale Characterization of Genomic Structural Variation in the Human Genome Reveals Adaptive and Biomedically Relevant Hotspots. Genome Biol. Evol. 11, 1136–1151 (2019).

13. Sedlazeck, F. J. et al. Accurate detection of complex structural variations using single-molecule sequencing. Nat. Methods 15, 461–468 (2018).

14. Sudmant, P. H. et al. An integrated map of structural variation in 2,504 human genomes. Nature 526, 75–81 (2015).

15. Collins, R. L. et al. A structural variation reference for medical and population genetics. Nature 581, 444–451 (2020).

16. He, Y. et al. De novo assembly of a Tibetan genome and identification of novel structural variants associated with high-altitude adaptation. Natl. Sci. Rev. 7, 391–402 (2020).

17. Sultana, T. et al. The Landscape of L1 Retrotransposons in the Human Genome Is Shaped by Pre-insertion Sequence Biases and Post-insertion Selection. Mol. Cell 74, 555–570.e7 (2019).

18. Weckselblatt, B. & Rudd, M. K. Human Structural Variation: Mechanisms of Chromosome Rearrangements. Trends Genet. 31, 587–599 (2015).

19. Xu, S. et al. A genome-wide search for signals of high-altitude adaptation in Tibetans. Mol. Biol. Evol. 28, 1003–1011 (2011).

20. Petousi, N. & Robbins, P. A. Human adaptation to the hypoxia of high altitude: the Tibetan paradigm from the pregenomic to the postgenomic era. J. Appl. Physiol. 116, 875–884 (2014).

21. Peng, Y. et al. Genetic variations in Tibetan populations and high-altitude adaptation at the Himalayas. Mol. Biol. Evol. 28, 1075–1081 (2011).

22. Szpiech, Z. A., Novak, T. E., Bailey, N. P. & Stevison, L. S. High-altitude adaptation in rhesus macaques. doi:10.1101/2020.05.19.104380.

23. Grotenboer, N. S., Ketelaar, M. E., Koppelman, G. H. & Nawijn, M. C. Decoding asthma: translating genetic variation in IL33 and IL1RL1 into disease pathophysiology. J. Allergy Clin. Immunol. 131, 856–865 (2013).

24. Wu, D.-D. et al. Convergent genomic signatures of high-altitude adaptation among domestic mammals. Natl Sci Rev 7, 952–963 (2019).

25. Deng, L. et al. Prioritizing natural-selection signals from the deep-sequencing genomic data suggests multi-variant adaptation in Tibetan highlanders. Natl Sci Rev 6, 1201–1222 (2019).

26. Boix, C. A., James, B. T., Park, Y. P., Meuleman, W. & Kellis, M. Regulatory genomic circuitry of human disease loci by integrative epigenomics. Nature (2021) doi:10.1038/s41586-020-03145-z.

27. Kang, R. et al. EnhancerDB: a resource of transcriptional regulation in the context of enhancers. Database 2019, (2019).

28. Chakraborty, R., Sikarwar, A. S., Hinton, M., Dakshinamurti, S. & Chelikani, P. Characterization of GPCR signaling in hypoxia. Methods Cell Biol. 142, 101–110 (2017).

29. Ding, D. et al. Genetic variation in PTPN1 contributes to metabolic adaptation to high-altitude hypoxia in Tibetan migratory locusts. Nat. Commun. 9, 4991 (2018).

30. Jia, Z. et al. Impacts of the Plateau Environment on the Gut Microbiota and Blood Clinical Indexes in Han and Tibetan Individuals. mSystems 5, (2020).

31. Eriksson, J. et al. Prolyl 4- hydroxylase subunit alpha 1 (P4HA1) is a biomarker of poor prognosis in primary melanomas, and its depletion inhibits melanoma cell invasion and disrupts tumor blood vessel walls. Molecular Oncology vol. 14 742–762 (2020).

32. Yun, C. et al. Proteasomal adaptation to environmental stress links resistance to proteotoxicity with longevity in Caenorhabditis elegans. Proc. Natl. Acad. Sci. U. S. A. 105, 7094–7099 (2008).

33. Liao, C. et al. Multi-tissue probabilistic fine-mapping of transcriptome-wide association study identifies cis-regulated genes for miserableness. doi:10.1101/682633.

34. Li, Y. et al. Hypoxia potentially promotes Tibetan longevity. Cell Res. 27, 302–305 (2017).

35. Zhou, Y. et al. Hypoxia augments LPS-induced inflammation and triggers high altitude cerebral edema in mice. Brain Behav. Immun. 64, 266–275 (2017).

36. Lachmann, A. et al. Geneshot: search engine for ranking genes from arbitrary text queries. Nucleic Acids Res. 47, W571–W577 (2019).

37. Jiang, C. et al. Chronic mountain sickness in Chinese Han males who migrated to the Qinghai-Tibetan plateau: application and evaluation of diagnostic criteria for chronic mountain sickness. BMC Public Health 14, 701 (2014).

38. Moore, L. G., Zamudio, S., Zhuang, J., Sun, S. & Droma, T. Oxygen transport in tibetan women during pregnancy at 3,658 m. Am. J. Phys. Anthropol. 114, 42–53 (2001).

39. Kanehisa, M., Furumichi, M., Tanabe, M., Sato, Y. & Morishima, K. KEGG: new perspectives on genomes, pathways, diseases and drugs. Nucleic Acids Res. 45, D353–D361 (2017).

40. James, C. et al. Herpes simplex virus: global infection prevalence and incidence estimates, 2016. Bull. World Health Organ. 98, 315–329 (2020).

41. The Gene Ontology Consortium. The Gene Ontology Resource: 20 years and still GOing strong. Nucleic Acids Res. 47, D330–D338 (2019).

42. Ihaka, R. & Gentleman, R. R: A Language for Data Analysis and Graphics. J. Comput. Graph. Stat. 5, 299–314 (1996).

43. 1000 Genomes Project Consortium et al. A global reference for human genetic variation. Nature 526, 68–74 (2015).

44. Purcell, S. et al. PLINK: a tool set for whole-genome association and population-based linkage analyses. Am. J. Hum. Genet. 81, 559–575 (2007).

45. Tang, H., Peng, J., Wang, P. & Risch, N. J. Estimation of individual admixture: analytical and study design considerations. Genet. Epidemiol. 28, 289–301 (2005).

46. Danecek, P. et al. The variant call format and VCFtools. Bioinformatics 27, 2156–2158 (2011).

47. Robinson, J. T. et al. Integrative genomics viewer. Nat. Biotechnol. 29, 24–26 (2011).

48. Meyer, M. et al. A high-coverage genome sequence from an archaic Denisovan individual. Science 338, 222–226 (2012).

49. Prüfer, K. et al. The complete genome sequence of a Neanderthal from the Altai Mountains. Nature 505, 43–49 (2014).

50. Prüfer, K. et al. A high-coverage Neandertal genome from Vindija Cave in Croatia. Science 358, 655–658 (2017).

51. Fu, Q. et al. Genome sequence of a 45,000-year-old modern human from western Siberia. Nature 514, 445–449 (2014).

52. Li, H. Aligning sequence reads, clone sequences and assembly contigs with BWA-MEM. arXiv[q-bio.GN] (2013).

53. Ye, K., Schulz, M. H., Long, Q., Apweiler, R. & Ning, Z. Pindel: a pattern growth approach to detect break points of large deletions and medium sized insertions from paired-end short reads. Bioinformatics 25, 2865–2871 (2009).

54. Rausch, T. et al. DELLY: structural variant discovery by integrated paired-end and split-read analysis. Bioinformatics 28, i333–i339 (2012).

55. Jeffares, D. C. et al. Transient structural variations have strong effects on quantitative traits and reproductive isolation in fission yeast. Nat. Commun. 8, 14061 (2017).

56. Zhao, H. et al. CrossMap: a versatile tool for coordinate conversion between genome assemblies. Bioinformatics 30, 1006–1007 (2014).

57. Yu, G., Wang, L.-G., Han, Y. & He, Q.-Y. clusterProfiler: an R package for comparing biological themes among gene clusters. OMICS 16, 284–287 (2012).

