## Supplemental Figures for "Structural variant selection for high-altitude adaptation using single-molecule long-read sequencing"

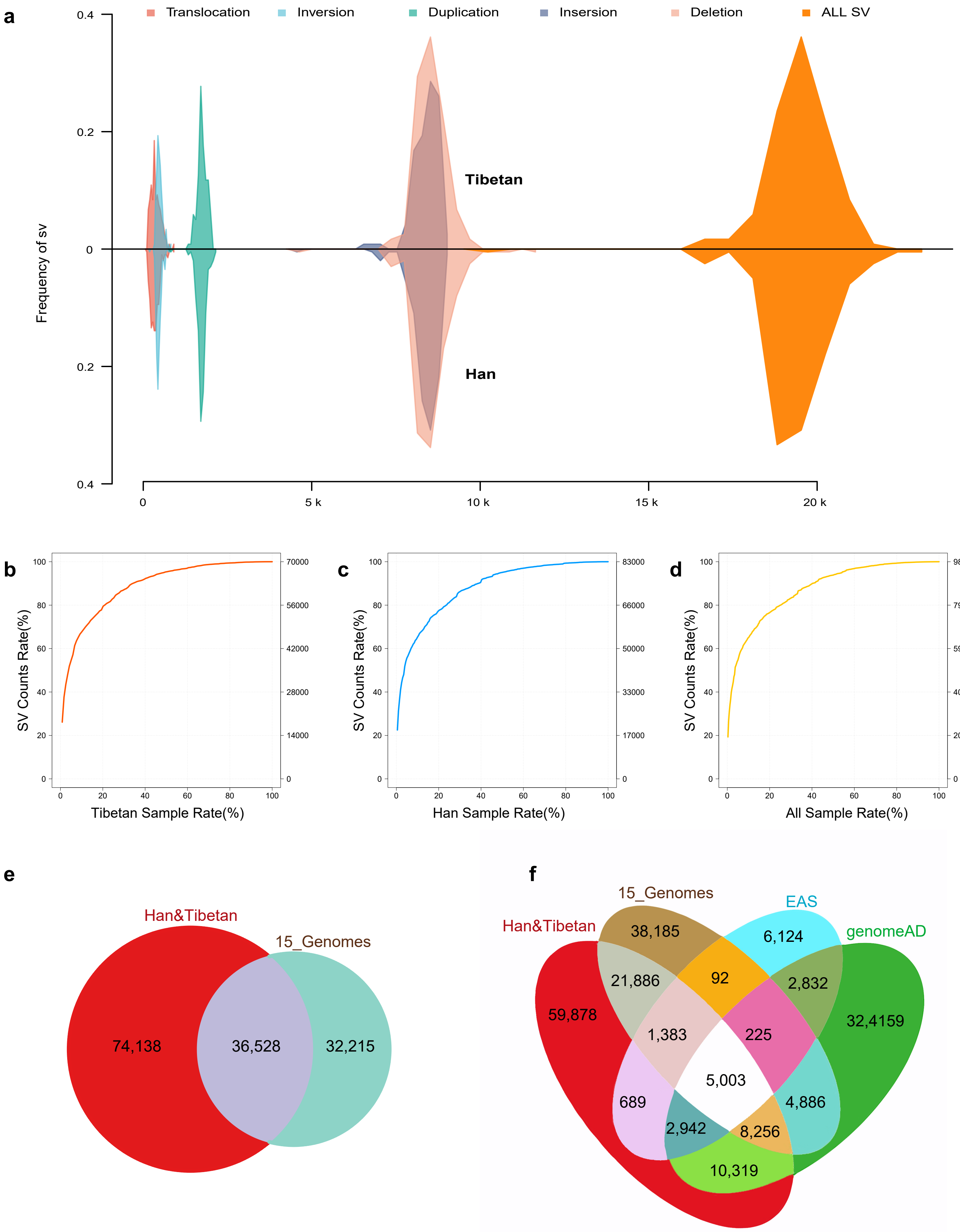


**Supplementary Figure 1. Han & Tibetan SV call set.**

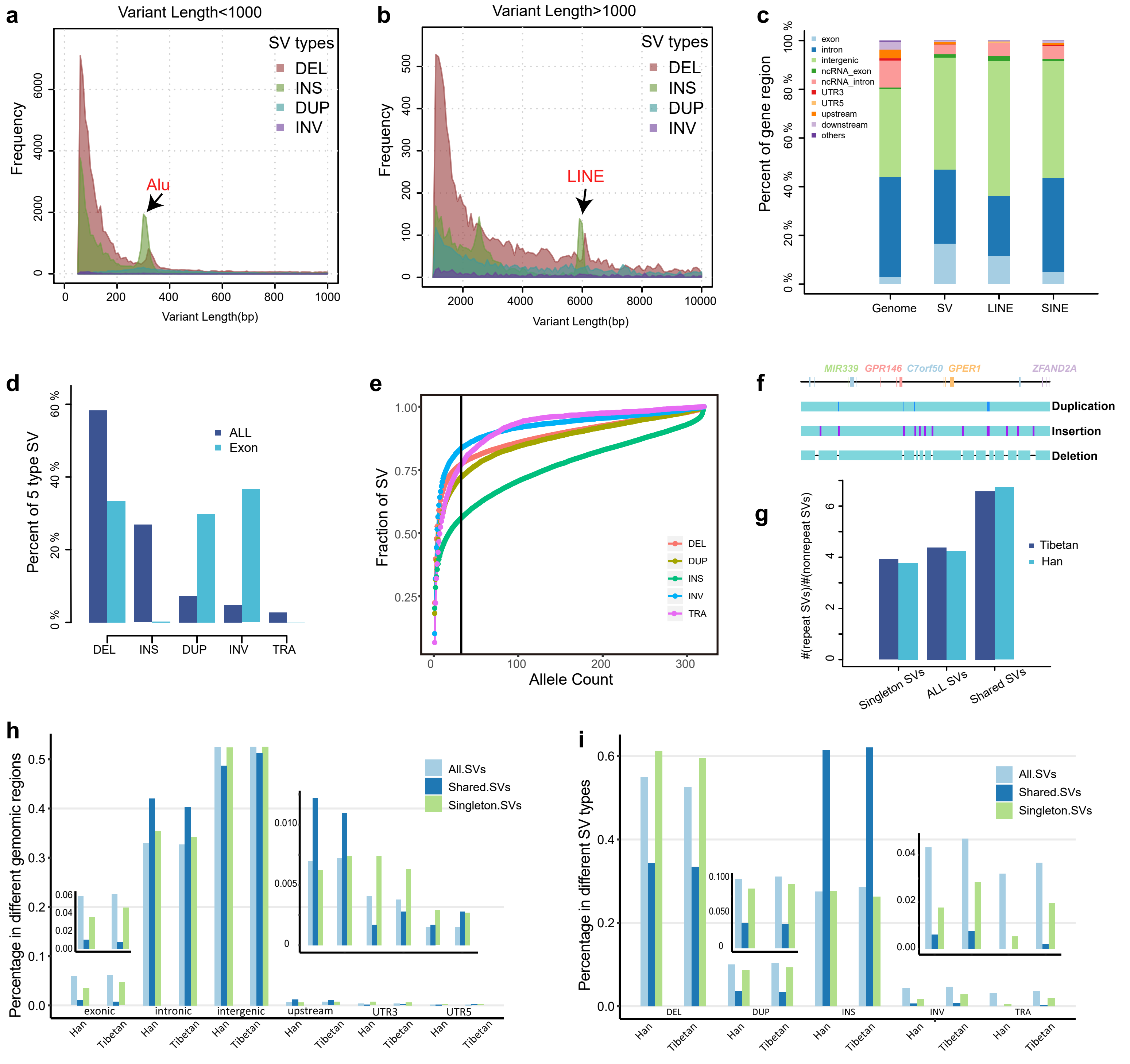


**Supplementary Figure 2. Characteristics of SV distribution and composition.**

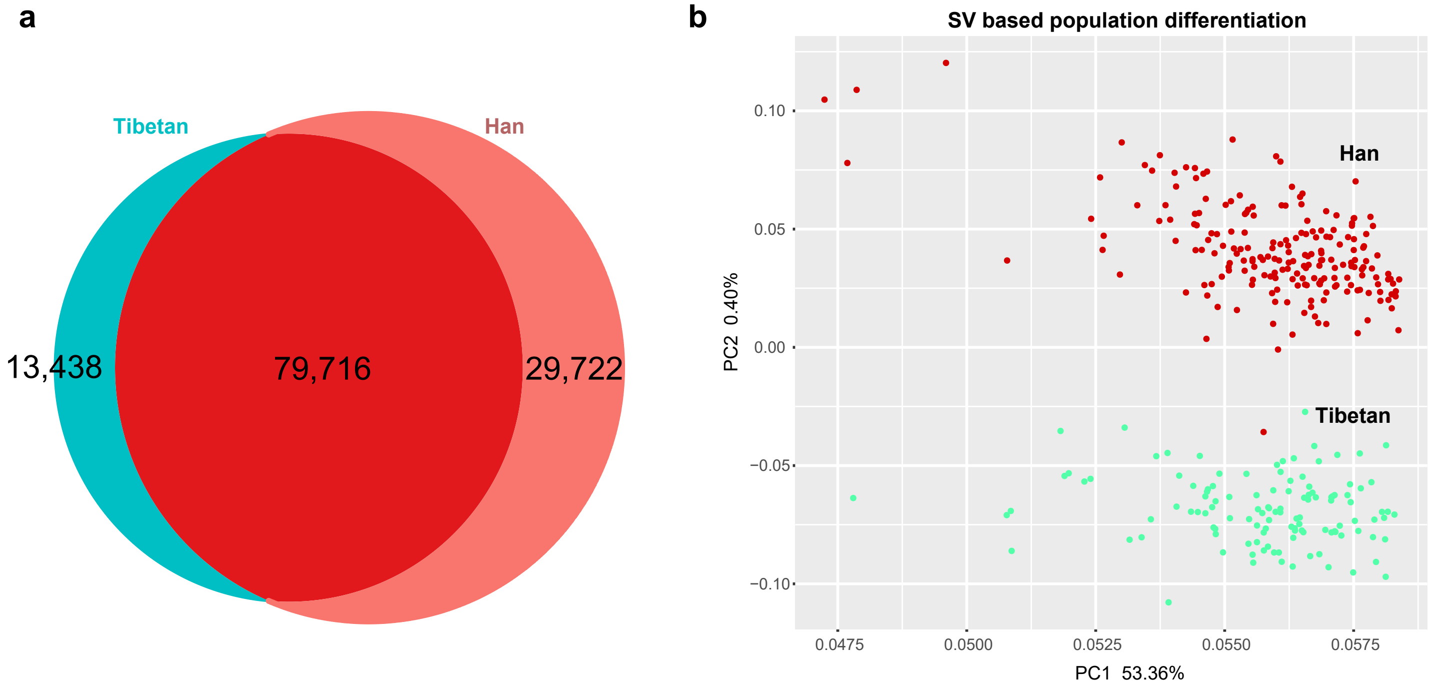


**Supplementary Figure 3. Population genetics of Han and Tibetan populations.**

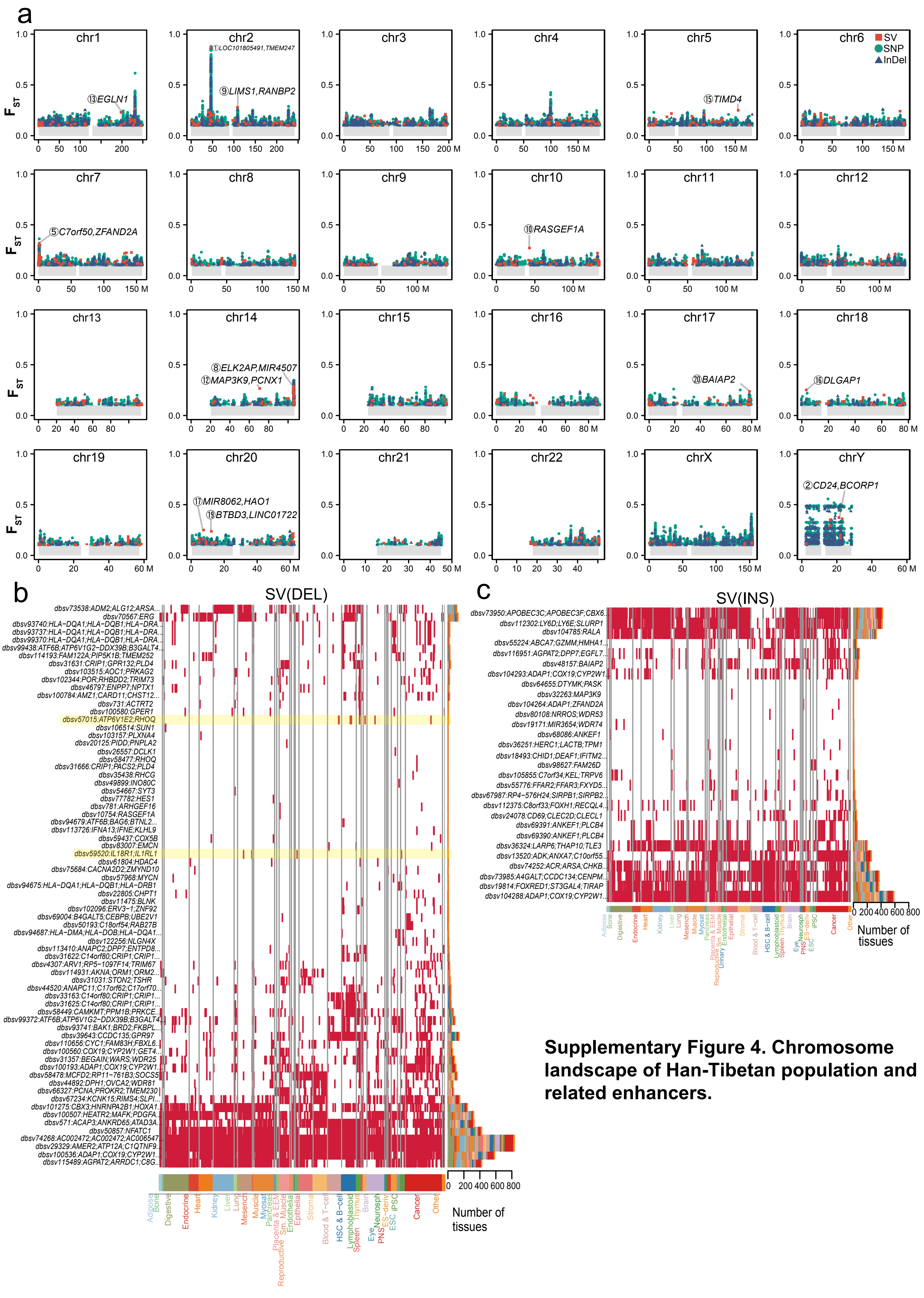


**Supplementary Figure 4. Chromosome landscape of Han-Tibetan population and related enhancers.**

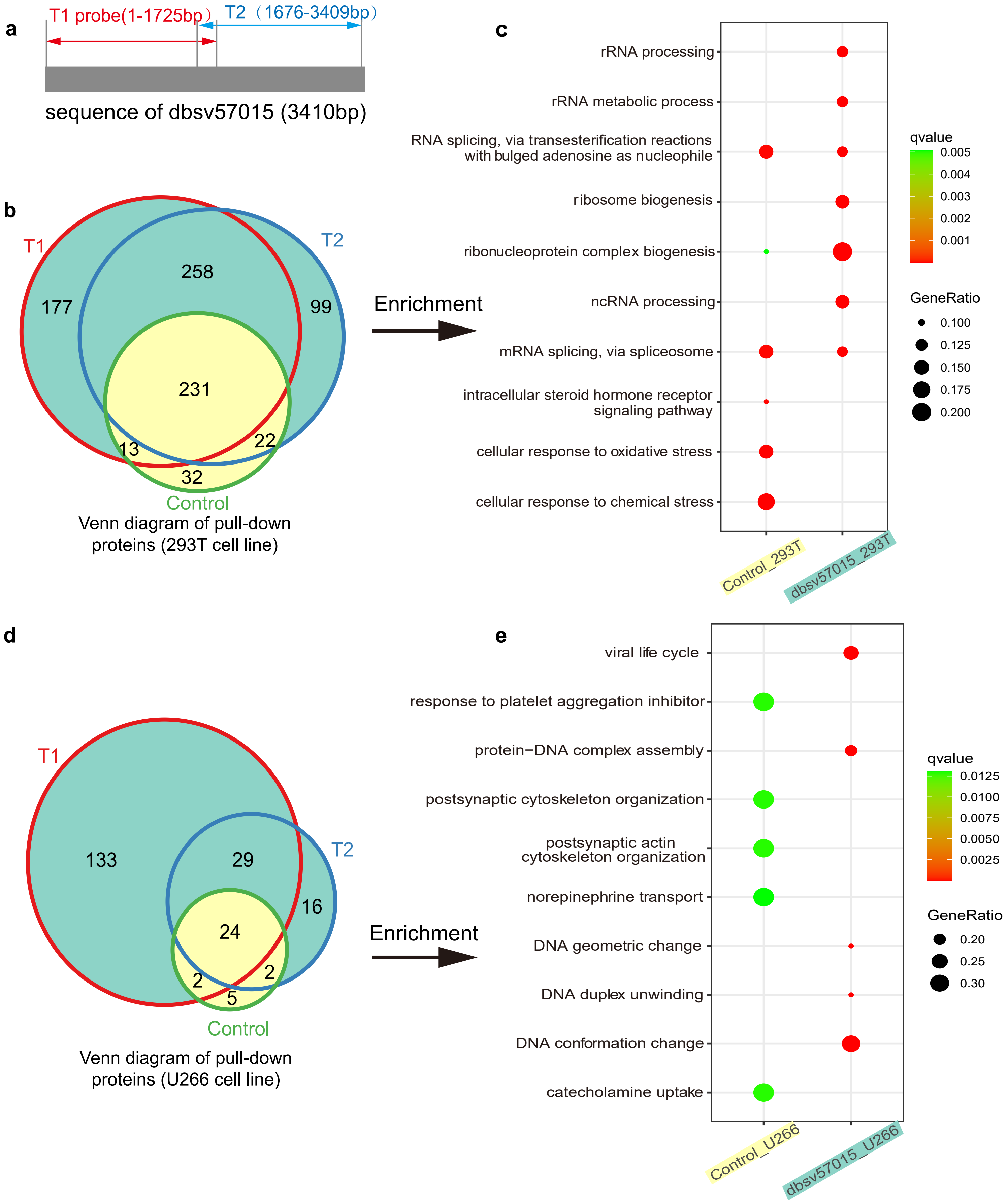


**Supplementary Figure 5. DNA pull-down results for the dbsv57015 sequence.**

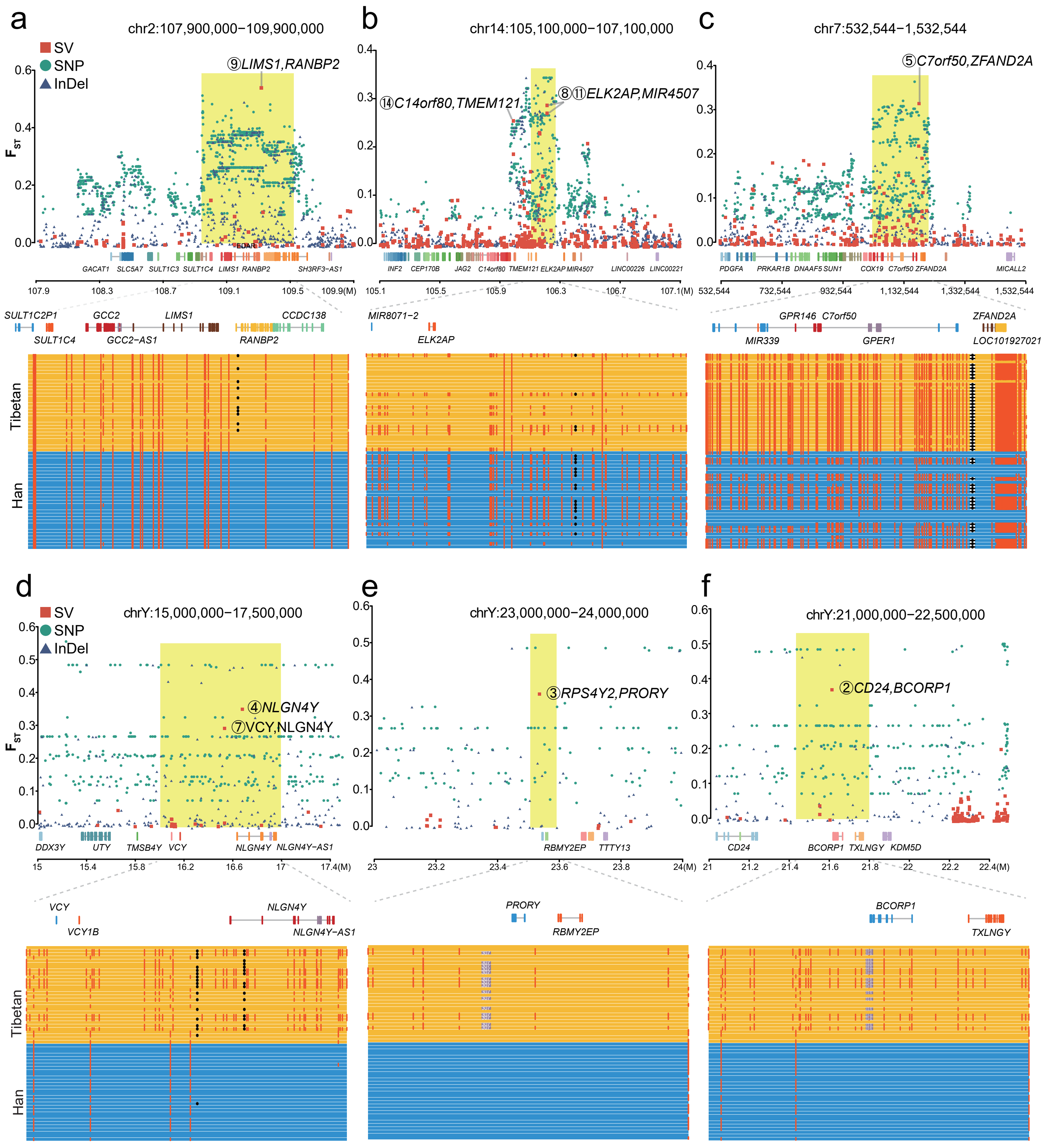


**Supplementary Figure 6. Manhattan plot of several population-specific regions.**

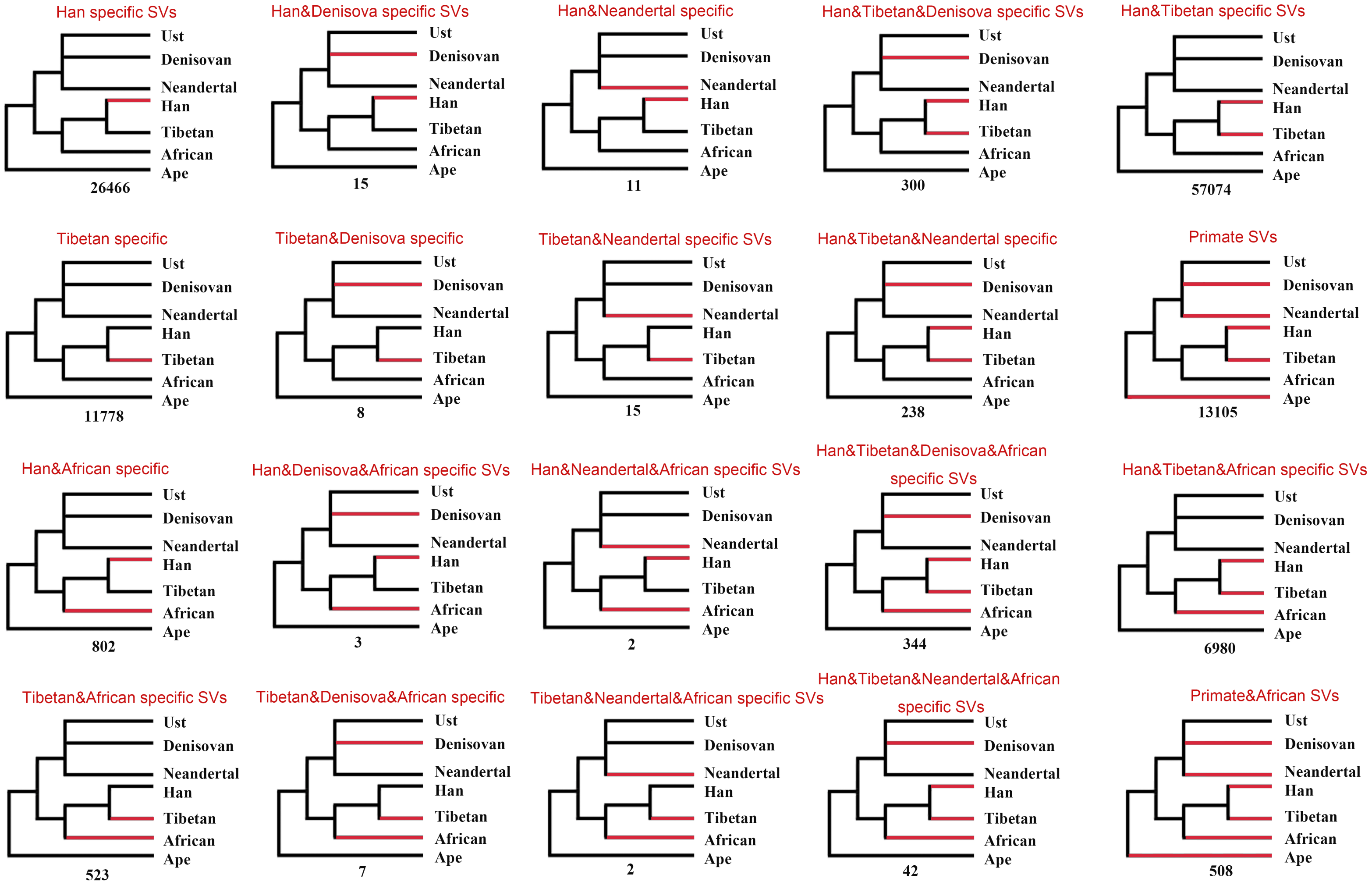


**Supplementary Figure 7. Possible evolutionary scenarios of SVs.**

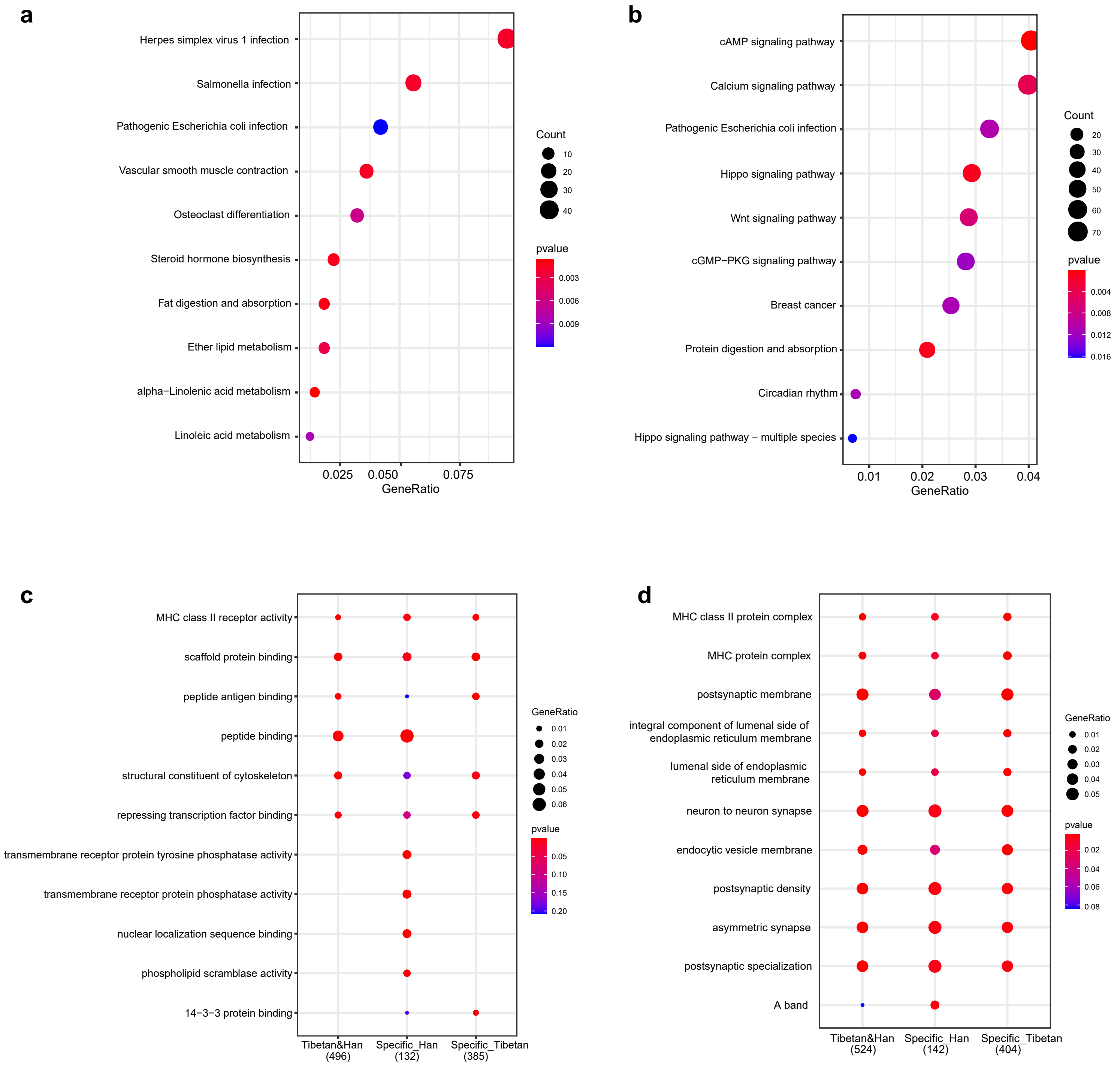


**Supplementary Figure 8. Enrichment analysis of Han/Tibetan-specific SVs.**
