## Supplemental Table 2-8 for "Structural variant selection for high-altitude adaptation using single-molecule long-read sequencing"

### Supplementary Tables

#### Supplementary Table 2. Statistics of ONT and SMRT-CCS sequencing data.

| **sample** | **num_of_reads* (M)** | **num_of_bases* (Gb)** | **fastq_depth (X)** | **Read length N50 (kb)** |
| --- | --- | --- | --- | --- |
| ONT data per sample | 3.82±1.53 | 62.55±22.62 | 20.85±7.54 | 22.71±4.04 |
| ONT data of AL-2-033 | 2.09 | 36.59 | 12.2 | 26.22 |
| SMRT PB data of AL-2-033 | 43.45 | 452.01 | 150.67 | 14.23 |

#### Supplementary Table 3. The orthogonal validation of ONT-SVs against SMRT-CCS-SVs by the same sample.

| **SV type** | **DEL** | **INS** | **INV** | **DUP** | **All** | **ratio** |
| --- | --- | --- | --- | --- | --- | --- |
| **SV number (CCS)** | 8,450 | 12,158 | 26 | 0 | 20,617 | 100% |
| **SV number(ONT, Guppy 2)** | 8,085 | 8,635 | 318 | 964 | 18,002 | 100% |
| **Common (CCS&Guppy 2)** | 5,659 | 8,738 | 19 | 0 | 14,416 | 70% |
| **Common (ONT, Guppy 2)** | 5,520 | 7,801 | 21 | 0 | 13,342 | 74% |
| **SV number(ONT, Guppy 3)** | 7,643 | 7,891 | 359 | 1,664 | 17,557 | 100% |
| **Common (CCS&Guppy 3)** | 5,546 | 7,822 | 20 | 0 | 13,388 | 65% |
| **Common (ONT, Guppy 3)** | 5,416 | 6,996 | 20 | 0 | 12,429 | 71% |

#### Supplementary Table 4. AF distribution for different types novel SV in Tibetan-Han population.

| **AF** | **DEL** | **INS** | **DUP** | **INV** | **Total** |
| --- | --- | --- | --- | --- | --- |
| **0**~**0.1** | 25,896 | 4,143 | 3,646 | 1,130 | 34,815 |
| **0.1~0.4** | 2,183 | 557 | 1,166 | 233 | 4,139 |
| **0.4~1** | 1,319 | 292 | 1,129 | 225 | 2,965 |
| **1** | 18 | 7 | 19 | 4 | 48 |
| **singleton** | 11,472 | 4,648 | 1,463 | 328 | 17,911 |
| **Total** | 40,888 | 9,647 | 7,423 | 1,920 | 59,878 |

#### Supplementary Table 5. SV comparison between ZF1 and our Tibetan population data

| **Type** | **DEL** | **DUP** | **INS** | **INV** | **TRA** | **ALL/Ratio** |
| --- | --- | --- | --- | --- | --- | --- |
| **Tibetan population** | 48,948 | 9,666 | 26,746 | 4,372 | 3,422 | 93154 (100%) |
| **ZF1** | 7,461 | 1,853 | 8,196 | 204 | 0 | 17714 (100%) |
| **Common（ZF1）** | 6,127 | 1,500 | 7,929 | 170 | 0 | 15726 (88.8%)（88.78%） |
| **Common（TP）** | 6,127 | 1,535 | 8,035 | 175 | 0 | 15890 (17.1%) |

#### Supplementary Table 6. Mean SV statistics for each sample of different AF in Tibetan-Han population

| **Allele frequency (AF)** | **SV number for each sample (Mean±SD）** | | | | | |
| --- | --- | --- | --- | --- | --- | --- |
|  | **Tibetan** | | | **Han** | | |
|  | **ALL** | **repeat region** | **non-repeat region** | **ALL** | **repeat region** | **non-repeat region** |
| **0~0.1** | 1,206.97±366.01 | 897.51±286.22 | 309.45±87.14 | 1,404.60±540.35 | 1,062.55±452.89 | 342.05±101.83 |
| **0.1~0.4** | 3,749.29±312.25 | 2,770.57±220.43 | 978.72±109.71 | 3,666.70±361.36 | 2,712.71±257.37 | 953.99±115.50 |
| **0.4~1** | 12,533.93±476.72 | 10,002.43±384.96 | 2,531.5±98.72 | 12,685.84±564.97 | 10,159.30±445.93 | 2,526.53±123.95 |
| **1** | 1,448±0 | 1,266±0 | 182±0 | 1,144±0 | 993±0 | 151±0 |
| **singleton** | 198.42±99.85 | 157.14±85.15 | 41.28±19.72 | 134.63±50.52 | 107.35±46.91 | 27.27±7.68 |

#### Supplementary Table 7. SV distribution in different genomic regions in Tibetan-Han population.

| **SV types** | **population** | **DEL (%)** | **INS (%)** | **DUP (%)** | **INV (%)** | **TRA (%)** | **Total (%)** |
| --- | --- | --- | --- | --- | --- | --- | --- |
| **LTR** | Tibetan | 2,972 (6.07) | 750（2.80） | 815 (8.43) | **848 (19.40)** | 220 (6.43) | 5,605 (6.02) |
|  | Han | 3,711 (6.17) | 819 （2.72） | 925 (8.44) | **910 (19.29)** | 230 (6.57) | 6,595 (6.03) |
| **Satellite** | Tibetan | 2,072 (4.23) | 273 （1.02） | 1,405 (14.54) | **745 (17.04)** | **708 (20.69)** | 5,203 (5.59) |
|  | Han | 2,367 (3.94) | 313 （1.04） | 1,519 (13.86) | **838 (17.76)** | **706 (20.15)** | 5,743 (5.25) |
| **Simple_repeat** | Tibetan | **10,635 (21.73)** | **8,186 (30.61) （30.61）** | **2,053 (21.24)** | 577 (13.20) | 77 (2.25) | **21,528 (23.11)** |
|  | Han | **12,744 (21.19)** | **8,844 (29.37)（29.37）** | **2,224 (20.29)** | 607 (12.87) | 87 (2.48) | **24,506 (22.39)** |
| **Segdup** | Tibetan | 1,832 (3.74) | 516 （1.93） | 413 (4.27) | 8 (0.18) | **1,177 (34.40)** | 3,946 (4.24) |
|  | Han | 2,163 (3.60) | 560 （1.86) | 473 (4.32) | 13 (0.28) | **1,243 (35.48)** | 4,452 (4.07) |
| **Other** | Tibetan | 2,929 (5.98) | 1,092 (4.08) | 445 (4.60) | 449 (10.27) | 84 (2.45) | 4,999 (5.37) |
|  | Han | 3,535 (5.88) | 1,334 (4.43) | 498 (4.54) | 478 (10.13) | 82 (2.34) | 5,927 (5.42) |
| **LINE** | Tibetan | **5,645 (11.53)** | **1,689 (6.32)** | **1,121 (11.60)** | **867 (19.83)** | **577 (16.86)** | **9,899 (10.63) (10.63)** |
|  | Han | **7,122 (11.84)** | **2,046 (6.80)** | **1,334 (12.17)** | **926 (19.63)** | **600 (17.13)** | **12,028 (10.99) (10.99)** |
| **SINE** | Tibetan | **11,381 (23.25)** | **6,473 (24.20)** | **1,058 (10.95)** | 681 (15.58) | 318 (9.29) | **19,911 (21.37) (21.37)** |
|  | Han | **14,656 (24.37)** | **7,704 (25.59)** | **1,327 (12.11)** | 732 (15.52) | 321 (9.16) | **24,740 (22.61) (22.61)** |
| **Low_complexity** | Tibetan | 2,692 (5.50) | 673 (2.52) | 753 (7.79) | 155 (3.55) | 19 (0.56) | 4,292 (4.61) |
|  | Han | 3,386 (5.63) | 717 (2.38) | 836 (7.63) | 172 (3.65) | 25 (0.71) | 5,136 (4.69) |
| **non-repeat region** | Tibetan | 8,790 (17.96) | 7,096 (26.53) | 1,603 (16.58) | 42 (0.96) | 242 (7.07) | **17,773 (19.08) (19.08)** |
|  | Han | 10,465 (17.40) | 7,774 (25.82) | 1,823 (16.63) | 42 (0.89) | 209 (5.97) | **20,313 (18.56) (18.56)** |
| **Total** | Tibetan | **48,948 (100)** | **26,746 (100)** | 9,666 (100) | 4,372 (100) | 3,422 (100) | 93,154 (100) |
|  | Han | **60,149 (100)** | **30,109 (100)** | 10,959 (100) | 4,718 (100) | 3,503 (100) | 109,438 (100) |

#### Supplementary Table 8. SINE and LINE associated SVs in various genomic functional regions.

| **Function regions** | **Population** | **Annotated with LINE** | | | | | | | | | | **Annotated with SINE** | | | | |
| --- | --- | --- | --- | --- | --- | --- | --- | --- | --- | --- | --- | --- | --- | --- | --- | --- |
|  |  | **DEL** | **INS** | **DUP** | **INV** | **TRA** | **ALL** | **Each / Total functional region(%)** | **DEL** | **INS** | **DUP** | | **INV** | **TRA** | **ALL** | **Each / Total functional region(%)** |
| exonic | Tibetan | 380 | 1 | 351 | 487 | 0 | 1219 | 12.31 | 333 | 4 | 294 | | 362 | 0 | 993 | 4.99 |
|  | Han | 448 | 1 | 406 | 508 | 0 | 1363 | 11.33 | 435 | 6 | 367 | | 390 | 0 | 1198 | 4.84 |
| intronic | Tibetan | 1558 | 471 | 236 | 33 | 54 | 2352 | 23.76 | 4857 | 2521 | 274 | | 32 | 29 | 7713 | 38.74 |
|  | Han | 2008 | 598 | 274 | 36 | 55 | 2971 | 24.7 | 6206 | 2992 | 331 | | 29 | 34 | 9592 | 38.77 |
| intergenic | Tibetan | 3266 | 1085 | 453 | 232 | 443 | 5479 | 55.35 | 5307 | 3391 | 410 | | 167 | 235 | 9510 | 47.76 |
|  | Han | 4125 | 1291 | 547 | 249 | 466 | 6678 | 55.52 | 6871 | 4031 | 521 | | 183 | 233 | 11839 | 47.85 |
| ncRNA_exonic | Tibetan | 82 | 3 | 36 | 97 | 3 | 221 | 2.23 | 87 | 16 | 35 | | 99 | 4 | 241 | 1.21 |
|  | Han | 91 | 3 | 49 | 106 | 4 | 253 | 2.1 | 104 | 15 | 42 | | 105 | 4 | 270 | 1.09 |
| ncRNA_intronic | Tibetan | 305 | 108 | 39 | 7 | 71 | 530 | 5.35 | 554 | 414 | 17 | | 8 | 44 | 1037 | 5.21 |
|  | Han | 386 | 131 | 48 | 14 | 68 | 647 | 5.38 | 704 | 488 | 31 | | 11 | 44 | 1278 | 5.17 |
| 3’-UTR | Tibetan | 8 | 4 | 0 | 1 | 2 | 15 | 0.15 | 36 | 36 | 2 | | 4 | 2 | 80 | 0.40 |
|  | Han | 9 | 6 | 1 | 1 | 2 | 19 | 0.16 | 51 | 48 | 3 | | 5 | 2 | 109 | 0.44 |
| 5’-UTR | Tibetan | 9 | 3 | 0 | 2 | 0 | 14 | 0.14 | 11 | 5 | 2 | | 2 | 0 | 20 | 0.10 |
|  | Han | 12 | 3 | 2 | 3 | 0 | 20 | 0.17 | 23 | 5 | 0 | | 2 | 0 | 30 | 0.12 |
| upstream | Tibetan | 17 | 8 | 2 | 4 | 1 | 32 | 0.32 | 61 | 37 | 14 | | 4 | 1 | 117 | 0.59 |
|  | Han | 17 | 7 | 1 | 3 | 2 | 30 | 0.25 | 93 | 42 | 19 | | 3 | 1 | 158 | 0.64 |
| downstream | Tibetan | 20 | 5 | 4 | 4 | 3 | 36 | 0.36 | 131 | 47 | 10 | | 3 | 3 | 194 | 0.97 |
|  | Han | 26 | 5 | 5 | 6 | 3 | 45 | 0.37 | 163 | 75 | 13 | | 4 | 3 | 258 | 1.04 |
| others | Tibetan | 0 | 1 | 0 | 0 | 0 | 1 | 0.01 | 4 | 2 | 0 | | 0 | 0 | 6 | 0.03 |
|  | Han | 0 | 1 | 1 | 0 | 0 | 2 | 0.02 | 6 | 2 | 0 | | 0 | 0 | 8 | 0.03 |
| ALL | Tibetan | 5645 | 1689 | 1121 | 867 | 577 | 9899 | - | 11381 | 6473 | 1058 | | 681 | 318 | 19911 | - |
|  | Han | 7122 | 2046 | 1334 | 926 | 600 | 12028 | - | 14656 | 7704 | 1327 | | 732 | 321 | 24740 | - |
